# The *Xanthomonas* type-III effector protein XopS stabilizes *Ca*WRKY40a to regulate defense hormone responses and preinvasion immunity in pepper (*Capsicum annuum*)

**DOI:** 10.1101/2021.03.31.437833

**Authors:** Margot Raffeiner, Suayib Üstün, Tiziana Guerra, Daniela Spinti, Maria Fitzner, Sophia Sonnewald, Susanne Baldermann, Frederik Börnke

## Abstract

A critical component of plant immunity against invading pathogens is the rapid transcriptional reprogramming of the attacked cell to minimize virulence. Many adapted plant bacterial pathogens use type III effector (T3E) proteins to interfere with plant defense responses, including the induction of immunity genes. The elucidation of effector function is essential to understanding bacterial pathogenesis. Here, we show that XopS, a T3E of *Xanthomonas campestris* pv. *vesicatoria* (*Xcv*), interacts with and inhibits the proteasomal degradation of the transcriptional regulator of defense gene expression WRKY40. Virus-induced gene silencing of *WRKY40* in pepper enhanced plant tolerance towards *Xcv* infection, indicating it represses immunity. Stabilization of WRKY40 by XopS reduces the expression of its targets including salicylic acid (SA)-responsive genes and the jasmonic acid (JA) signaling repressor *JAZ8*. *Xcv* bacteria lacking XopS display significantly reduced virulence when surface inoculated onto susceptible pepper leaves. XopS delivery by *Xcv*, as well as ectopic expression of XopS in Arabidopsis or *Nicotiana benthamiana* prevented stomatal closure in response to bacteria and biotic elicitors in a WRKY40 dependent manner. This suggests that XopS interferes with preinvasion as well as with apoplastic defense by manipulating WRKY40 stability and gene expression eventually altering phytohormone crosstalk to promote pathogen proliferation.

## Introduction

Plants employ a two-tiered immune system to detect and resist attempted invasion of microbial pathogens. In the first layer, cell-surface localized pattern-recognition receptors (PRRs) recognize conserved pathogen/microbe-associated molecular patterns (PAMPs/MAMPs) and initiate PAMP/pattern-triggered immunity (PTI) (Zipfel, 2014; Macho and Zipfel, 2015). MAMP recognition induces a diverse array of defense responses, including the production of reactive oxygen species (ROS), cellular Ca^2+^ spikes, the activation of mitogen-activated protein kinases (MAPKs), synthesis of defense hormones such as salicylic acid (SA), and callose deposition at the cell wall (Boller and Felix, 2009). Collectively these events lead, in a largely unknown manner, to inhibition of pathogen growth and suppression of disease. Due to the conserved nature of MAMPs and the overlapping responses triggered by MAMPs derived from different organisms, PTI is effective against a broad range of non-adapted microbes. In contrast, adapted pathogens have acquired a range of virulence mechanisms to suppress PTI below a threshold of effective resistance and are thus able to proliferate and to cause disease on a particular host. Virulence mechanisms include the delivery of so called effector proteins directly into the host cell where these interfere with PTI responses or otherwise help the pathogen to establish a beneficial environment, eventually resulting in effector-triggered susceptibility (ETS) (Dou and Zhou, 2012; Langin et al., 2020; Macho and Zipfel, 2015). As a counter measure, plants have evolved a second layer of immunity which enables direct or indirect recognition of such effectors through cognate plant disease resistance (*R)* proteins, resulting in the activation of so-called effector-triggered immunity (ETI) (Dodds and Rathjen, 2010; Cesari, 2018). Generally, PTI and ETI give rise to similar responses, although ETI is qualitatively stronger and faster and often involves a rapid form of localized cell death called the hypersensitive response (HR) that is assumed to limit spread of biotrophic pathogens from the site of infection (Tsuda and Katagiri, 2010).

Pathogen invasion induces large scale reprogramming of plant gene expression, which is essential for launching a robust defense response (Tsuda and Somssich, 2015; Garner et al., 2016; Li et al., 2016). Infection of plants with a nonpathogenic *Pseudomonas syringae* pv. tomato (*Pst*) strain or treatment with the *Pst* derived MAMP flagellin 22 (flg22), identified 4000 MAMP-responsive genes in Arabidopsis (Lewis et al., 2015) and in tomato (Rosli et al., 2013). The genes induced early during PTI are related to defense responses and salicylic acid (SA) biosynthesis, whereas genes associated with photosynthesis and chloroplast functions are suppressed, suggesting that plants may actively reduce the production of photosynthates to restrict resource allocation for pathogen growth (Rosli et al., 2013; Lewis et al., 2015). Transcriptional reprogramming during induced defense is governed by interacting networks of transcription factors (TFs) from multiple families and their co-regulatory proteins which affect TF function through various molecular mechanisms (Moore et al., 2011). In general, these TFs often act downstream of MAPK cascades or Ca^2+^ signaling via diverse activation mechanisms to relay MAMP signaling into an appropriate transcriptional response. Prominent among the TF genes that were shown to mediate transcriptional reprogramming induced during PTI in Arabidopsis and other plant species, are members of the WRKY family of TFs (Pandey and Somssich, 2009; Tsuda and Somssich, 2015; Birkenbihl et al., 2017). Common to all WRKY TFs is the app. 60 amino acid long WRKY domain that binds to the W-box cis-element ((T)TGAC(C/T)) within the promoter of their target genes. Indeed, putative W-boxes have been identified in the promoter regions of many genes associated with plant biotic stress responses, including those involved in SA biosynthesis, and WRKYs have been described as both transcriptional activators and repressors in these pathways. In addition, WRKYs often form positive feedback regulatory loops via binding to their own promoters (Mao et al., 2011). Regulation of WRKY transcriptional activity also occurs at the post-translational level through phosphorylation or proteasomal protein turnover (Ishihama and Yoshioka, 2012; Matsushita et al., 2013).

In Arabidopsis, the closely related WRKY TFs WRKY18, and −40 have partially redundant functions in negatively regulating resistance to *P. syringae* and the biotrophic fungus *Golovinomyces orontii* (Xu et al., 2006; Pandey et al., 2010). *wrky40 wrky18* double mutants were more resistant towards both pathogens and displayed massive transcriptional reprogramming during early stages of infection in Arabidopsis (Pandey et al., 2010). Expression of several positive and negative regulators of jasmonic acid (JA) and SA signaling was significantly altered in resistant *wrky18 wrky40* double mutants compared with susceptible wild-type plants. About 1400 possible target genes for WRKY40 in Arabidopsis have been identified by ChIP-seq, with an enrichment for genes involved in the early processes of MAMP perception and signaling (Birkenbihl et al., 2017).

Bacterial type III effector proteins (T3Es) target immunity components at seemingly every stage of plant defense, including host transcriptional regulation (Macho and Zipfel, 2015; Büttner, 2016; Khan et al., 2018). T3Es can affect defense gene expression either indirectly by interfering with upstream phosphorelay signaling or hormonal cues, or through direct interaction with components of the transcriptional machinery and its auxiliary proteins. For example, the *Pst* T3E AvrPtoB targets a number of PRRs at the plasma membrane and mediates their proteasomal degradation by virtue of its E3-ubiquitin ligase activity (Göhre et al., 2008; Gimenez-Ibanez et al., 2009). AvrPtoB mediated inactivation of PRRs disrupts PAMP signaling and attenuates induction of defense genes. In addition, AvrPtoB also mediates degradation of NPR1, a transcriptional co-regulator in SA-dependent immunity, and thus directly interferes with SA-mediated gene expression (Chen et al., 2017).

The Gram-negative bacterium *Xanthomonas campestris* pv. *vesicatoria* (*Xcv*; synonymously designated as *Xanthomonas euvesicatoria*) is the causal agent of bacterial spot disease in pepper and tomato plants (Jones et al., 1998). Like for other foliar bacterial pathogens, a successful *Xcv* disease cycle proceeds from an epiphytic phase upon arrival on the surface of a healthy leaf to a largely endophytic phase accompanied by aggressive bacterial multiplication within the apoplast of infected tissues (McGuire et al., 1991). Invasion of the apoplast occurs through natural openings in the leaf surface, including stomata and hydatodes, and through wounds (Ramos and Volin, 1987). It has been shown that stomata play an active role in pre-invasion immunity against bacteria (Melotto et al., 2006). Sensing of MAMPs by PRRs on the surface of guard cells around the stomatal pore induces stomatal closure preventing further ingress of bacteria into the apoplast, a mechanism referred to as stomatal immunity (Melotto et al., 2006; Sawinski et al., 2013). In addition to PAMP perception, the plant hormones abscisic acid (ABA), SA, and JA-Ile play integral roles in regulating stomatal immunity. Treatment of tomato plants with ABA to induce stomatal closure prior to surface inoculation with *Xcv* reduced disease incidence and severity of symptom development (Ramos and Volin, 1987), indicating that stomatal aperture represents a major limiting factor for disease progression. Several adapted bacterial pathogens have evolved secreted virulence factors to overcome stomatal immunity, including phytotoxins and T3Es that are able to open the stomatal pore (Melotto et al., 2006; Gimenez-Ibanez et al., 2014; Hurley et al., 2014). The best characterized example is the small molecule coronatine (COR) produced by *Pst* DC3000 to reopen closed stomata, thereby significantly increasing the number of entry sites for bacterial invasion.

COR is a structural mimic of JA-Ile (Krumm et al., 1995; Staswick and Tiryaki, 2004; Melotto et al., 2006; Okada et al., 2009) binding to the JA co-receptor COI1 (Katsir et al., 2008). When COR binds to COI1, downstream signaling leads to the induction of NAC transcription factors that repress SA biosynthesis genes and induce SA metabolism genes, thereby suppressing SA accumulation and promoting stomatal opening (Zheng et al., 2012; Du et al., 2014; Gimenez-Ibanez et al., 2017). *Xcv* is not known to produce COR, nor have other virulence factors targeting stomatal immunity of host plants been described for this pathogen. Thus, whether and how *Xcv* interferes with preinvasive defense responses remains elusive. The ability of *Xcv* to cause disease is largely dependent on a suite of app. 35 T3Es several of which are conserved between different *Xcv* strains or even *Xanthomonas* species and constitute a so called “core” set of effectors, while others are only contained in certain strains (Potnis et al., 2011; Schwartz et al., 2015). While T3Es from *Pseudomonas* appear to target predominantly the host cell plasma membrane, a large portion of *Xanthomonas* effector proteins are predicted to locate the plant cell nucleus (Khan et al., 2018). Although cellular targets and modes of action are known for only a limited number of *Xcv* effectors, it appears that direct manipulation of host transcription constitutes one of the key mechanisms underpinning bacterial pathogenesis (Boch and Bonas, 2010). Certain *Xcv* strains possess transcription activator-like (TAL) effectors, which mimic eukaryotic TFs by directly inducing host transcriptional reprogramming. For example, the TAL effector AvrBs3 activates the transcription of the basic helix-loop-helix (bHLH) TF UPA20 in pepper leaves to regulate cell size (Kay et al., 2007). XopD, another *Xcv* effector, promotes bacterial growth and delays symptom development in infected tomato plants (Kim et al., 2008). It possesses a C-terminal cysteine protease activity exhibiting small ubiquitin-like modifier (SUMO)-specific protease activity, two EAR (ERF-associated Amphiphilic Repression) transcriptional repressor motifs and a bHLH domain required for DNA binding and nuclear localization (Kim et al., 2008; Kim et al., 2013). XopD represses ethylene stimulated defense gene transcription in tomato by targeting and de-sumoylating the TF ERF4 (ethylene response factor 4). Removal of SUMO from ERF4 results in its destabilization by the proteasome (Kim et al., 2008; Kim et al., 2013).

An additional *Xcv* translocated T3E is XopS, which has originally been identified in the genome of *Xcv* strain 85-10 (Schulze et al., 2012; Teper et al., 2016). XopS is a protein of app. 34 kDa confined to the Xanthomonads sharing no obvious sequence similarity with other proteins found in the database. Thus, the lack of discernible structural features in XopS has yet precluded any prediction either of its function or its biochemical activity. An *XcvΔxopS* deletion strain displays reduced symptom development upon pressure infection of susceptible pepper plants although lack of the effector protein does not appear to affect bacterial multiplication in infected tissue (Schulze et al., 2012). When expressed in Arabidopsis protoplasts XopS interferes with the induction of PTI marker gene expression, likely acting downstream of MAP kinase signaling (Schulze et al., 2012; Popov et al., 2016). However, its host target proteins are unknown.

In the present study, we show that XopS interferes with pre-invasive as well with post-invasive defense responses and that it represents a major virulence factor to overcome stomatal immunity in a compatible interaction of *Xcv* with pepper plants. XopS interacts with the negative regulator of defense gene transcription WRKY40 inside the plant cell nucleus resulting in dampening of SA-dependent gene expression in favor of JA-mediated responses. XopS binding to WRKY40 stabilizes the TF through inhibition of its proteasomal degradation and thereby perpetuates its repressor activity during defense to eventually attenuate induction of WRKY40 target genes.

## Results

### XopS is a bacterial virulence factor and suppresses plant defense

Since susceptible pepper plants have reduced symptom development after infection with a *xopS* deletion strain (Schulze et al., 2012), we used an independently constructed *Xcv* 85-10 *xopS* mutant to re-investigae symptom development after infection of pepper leaves. Our results confirmed that the *Xcv*Δ*xopS* strain produced less symptoms between 4 to 6 days post inoculation (dpi) in leaves of susceptible pepper ECW plants as compared to wild type *Xcv* infected tissue (Supplementary Figure S1). While *Xcv* wild type infected leaves showed tissue collapse and clear signs of necrotic lesions, *Xcv*Δ*xopS* infected tissue displayed only slight chlorosis. Infection with an *Xcv*Δ*xopS(XopS-HA)* complementation strain restored the wild type phenotype (Supplementary Figure S1). *Xcv*Δ*xopS* infected plants displayed significantly higher levels of the defense hormone SA relative to *Xcv* wild type inoculated plants, which was accompanied by elevated expression of the SA-responsive defense gene *PR1* (Supplementary Figure S2A and B). This indicates that XopS interferes with cellular processes that lead to the development of host tissue necrosis and likely interferes with defense hormone balance.

*In planta* growth of an *Xcv*Δ*xopS* strain was not affected when bacteria were pressure infiltrated into leaves of susceptible pepper plants (Schulze et al., 2012), suggesting only a minor contribution of XopS to overall bacterial virulence under these experimental conditions. Pressure infiltration of pathogenic bacteria into susceptible leaf tissue circumvents the stomatal defense system as a major barrier of the plant immune system that bacteria encounter in a natural infection situation (Melotto et al., 2008). Adapted pathogens evolved several virulence factors including T3Es to overcome stomatal immunity in their respective hosts (Melotto et al., 2008; Hurley et al., 2014). In order to investigate whether XopS might affect pre-invasion defense responses, susceptible ECW pepper plants where dip-inoculated onto the surface of plant leaves (1×10^8^ CFU⋅ml^−1^). *Xcv*Δ*xopS* bacteria achieved significantly lower population densities than wild type bacteria at 7 dpi (Figure 1A). Complementation of the *Xcv*Δ*xopS* strain by ectopic XopS expression fully restored wild type growth. As observed upon pressure infiltration, dip-inoculation with the *Xcv*Δ*xopS* strain was accompanied by less severe symptom development retaining higher chlorophyll levels when compared to *Xcv* wild type or *Xcv*Δ*xopS(XopS-HA)* infected leaves (Figure 1B and C). Incubation of pepper leaves with *Xcv* wild type bacteria for two hours led to increased stomatal aperture as compared to mock treated leaves (Figure 1D). However, treatment of leaves with the *Xcv*Δ*xopS* strain or with a *XcvΔhrpF* strain, which is unable to deliver T3Es into the host cell, induced significant stomatal closure, while the *Xcv*Δ*xopS(xopS-HA)* strain was able to maintain stomatal opening. These results suggest that *Xcv* is able to increase stomatal aperture in a T3E dependent manner and that XopS is a major effector contributing to this effect. Similarly, XopS was able to induce stomatal opening when transiently expressed in leaves of *N. benthamiana* and XopS expression significantly inhibited stomatal closure in response to the treatment of leaves with the bacterial MAMP flg22, but did not affect the stomatal response to abscisic acid (ABA) (Supplementary Figure S3A and B), indicating that in these plants stomatal closure can occur normally upon cues other than MAMP perception. Expression of both transiently expressed proteins, GFP and XopS-GFP, was verified by western blotting (Supplementary Figure S3C).Taken together, our data demonstrate that XopS interferes with early defense responses, likely including stomatal immunity and thus contributes to bacterial virulence in a compatible *Xcv* host interaction.

**Figure 1.**
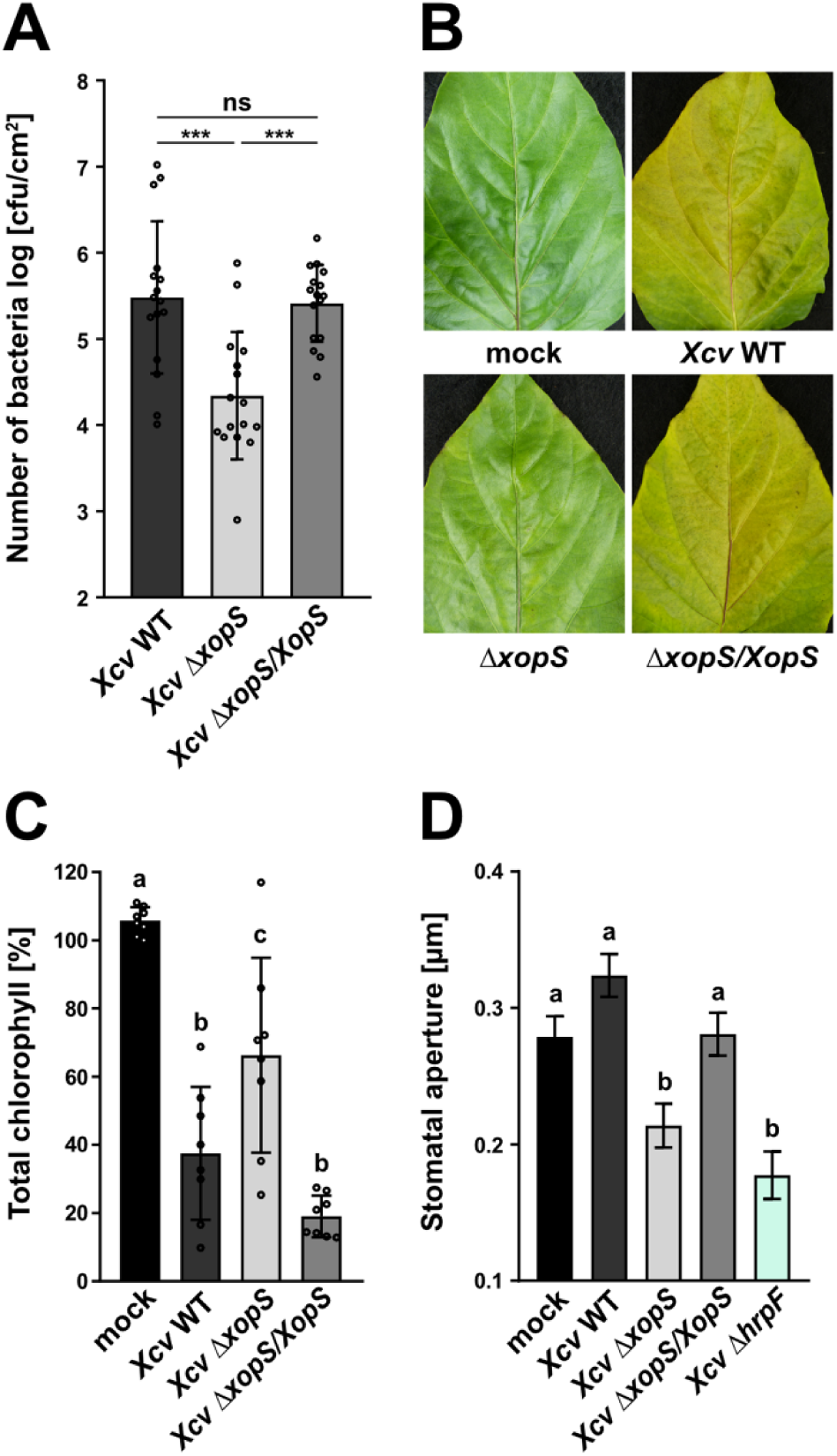
XopS is required for full virulence of *Xcv* on susceptible pepper plants. **(A)** Bacterial multiplication of *Xcv* wild type, *XcvΔxopS* and *XcvΔxopS(XopS-HA)* strains in susceptible pepper plants. Leaves were dip-inoculated with a bacterial suspension at OD_600_ = 0.2 and colony forming units (CFU) in infected tissue were quantified at 7 days post inoculation (dpi). Bars represent the mean of n = 8 biological replicates (and 2 technical replicates per biological replicate) ± SD. Significant differences are marked by asterisks (***, P < 0.001; ns, not significant) according to one-way ANOVA. The experiment was carried out three times with similar results. **(B)** Disease symptom development in pepper leaves, dip-inoculated with *Xcv* wild type, *XcvΔxopS* or *XcvΔxopS(XopS-HA)* strains. Pictures were taken at 7 dpi. A similar phenotype was observed every time plants were dip-inoculated with indicated strains. **(C)** Chlorophyll content in pepper leaves, dip-inoculated with indicated *Xcv* strains. Uninfected leaves were used as mock control and the percentage of total chlorophyll was determined when compared to the mock control (set to 100%). The measurement was performed at 7 dpi. Bars represent the mean of n = 8 biological replicates ± SD. Letters over bars represent statistical significance determined by one-way ANOVA (P < 0.05). The experiment was carried out twice with similar results. **(D)** Stomatal apertures of leaf discs from pepper plants floated on water alone (mock) or on water containing *Xcv* wild type, *XcvΔxopS*, *XcvΔxopS(XopS-HA) or XcvΔhrpF* strains at an OD_600_ = 0.2. The measurement of stomatal aperture was performed 2 h post treatment. Approximately 100 apertures from n = 4 different plants were measured per individual treatment and are represented as width/length ratio. Bars represent the mean ± SE and letters over bars represent statistical significance determined by one-way ANOVA (P < 0.05). The experiment was carried out twice with similar results.

To investigate whether XopS interferes with PTI in general, transgenic Arabidopsis lines expressing a XopS-GFP fusion protein under control of a β-estradiol inducible promoter were generated and expression of the fusion protein was verified by immunoblotting (Supplementary Figure S4A). Independent transgenic Arabidopsis lines supported enhanced bacterial growth of a non-pathogenic *ΔhrcC Pseudomonas syringae* DC3000 strain upon induction of XopS-GFP expression (Supplementary Figure S4B). This indicates that XopS-GFP interferes with post-invasive PTI responses in Arabidopsis. Interestingly, the two transgenic lines with the highest XopS-GFP expression level (line 4 and 5) developed chlorotic symptoms at approximately 5 days after β-estradiol treatment, independent of bacterial inoculation (Supplementary Figure S5A). This phenotype is reminiscent of T3E expressing plants with an activated jasmonic acid (JA)-response (Gimenez-Ibanez et al., 2014). Thus, we analyzed expression of JA marker genes induced early in the JA response in a representative transgenic line. As shown in Supplementary Figure S5B, expression of JA marker genes *AtMYC2* and *AtJAZ10* was drastically induced in XopS-GFP plants 4 h after estradiol treatment. This is in line with elevated JA levels in XopS-GFP expressing Arabidopsis plants 24 h after induction with β-estradiol (Supplementary Figure S5C). In accordance with our previous observations, inducible expression of XopS-GFP in Arabidopsis prevented stomatal closure in response to treatment with flg22 (Supplementary Figure S5D).

Taken together, the experiments show that XopS interferes with PTI, MAMP-induced stomatal closure, and triggers a JA-response when ectopically expressed in Arabidopsis.

### XopS interacts with WRKY40 in yeast

To better understand how XopS manipulates host cellular processes, we screened for proteins that interact with XopS using a yeast two-hybrid (Y2H) cDNA library from tobacco (*Nicotiana tabacum*). Although this species is not a natural host plant for *Xcv*, our previous research suggests that the conservation of potential *Xcv* T3E target proteins between tobacco and pepper is sufficiently high to identify *bona fide* XopS host targets (Üstün et al., 2013a). One cDNA identified in the Y2H screening for XopS interacting proteins encoded a probable WRKY TF. WRKYs are one of the largest families of TFs in plants and, according to their domain structure, can be divided into three different groups (Rushton et al., 2010). Based on the polypeptide sequence, the WRKY TF identified in our screen belongs to group IIa and, when compared to the WRKY family of Arabidopsis as the best studied reference, it shows highest identity (ca. 64%) to Arabidopsis WRKY40 (Supplementary Figure S6). Thus, it was termed *Nt*WRKY40.

In Arabidopsis, WRKY40 together with other WRKY TFs acts as a negative regulator of defense against biotrophic pathogens (Xu et al., 2006; Pandey and Somssich, 2009) and has more recently been shown to negatively regulate PAMP-induced gene expression (Birkenbihl et al., 2017). In order to better understand the interaction of XopS with WRKY40, we tested the ability of the T3E to bind WKRY40 orthologs from other species including Arabidopsis, *N. benthamiana* and the *Xcv* host plant pepper. A pepper WRKY40 has previously been described to be involved in the positive regulation of both high temperature tolerance and *Ralstonia solanacearum* resistance (Dang et al., 2013). However, the pepper WRKY we investigate here (Ca03g32070) shows only 71% similarity to the previously described pepper WRKY40 (Ca00g87690) in a BLAST comparison, despite both WRKYs having highest similarity to WRKY40 when compared to the WRKY family from Arabidopsis (Supplementary Figure S6). In order to avoid confusion with the pepper WRKY40 described by Dang et al., (2013), we tentatively named the pepper WRKY analyzed for its ability to interact with XopS *Ca*WRKY40a. The direct interaction assays in yeast revealed that XopS was able to interact with WRKY40 proteins from Arabidopsis and *N. benthamiana* as well as with pepper WRKY40a (Figure 2).

**Figure 2.**
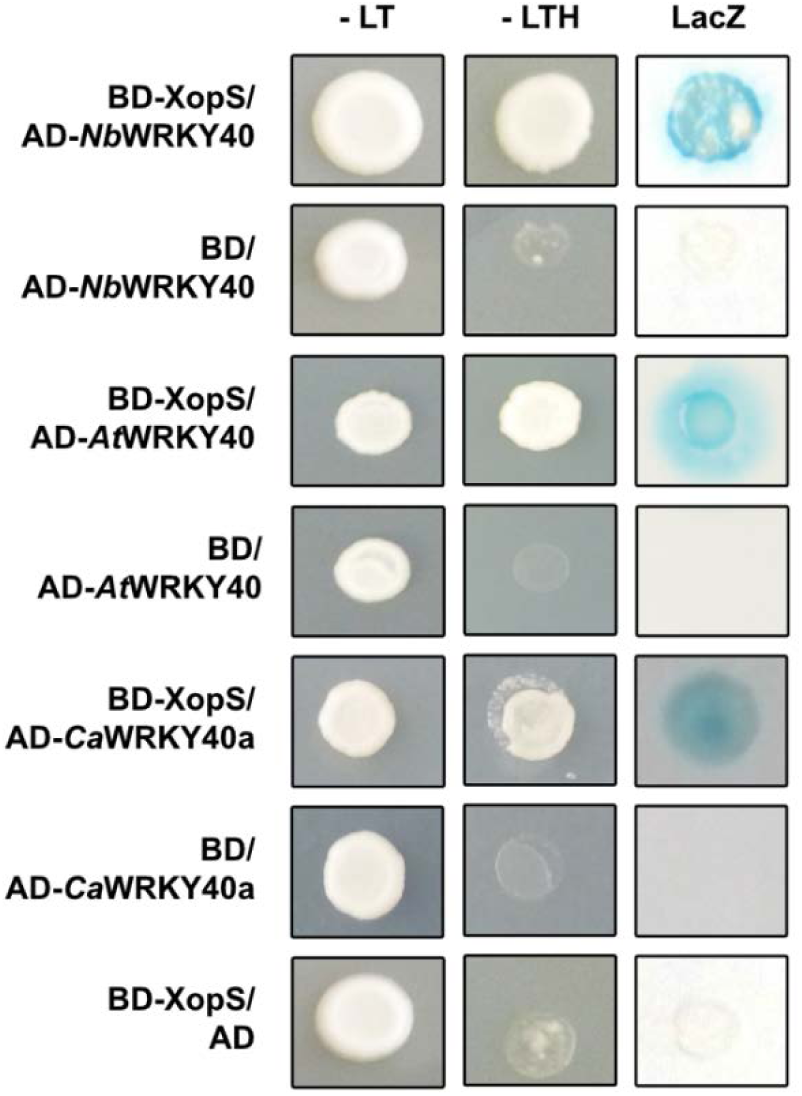
XopS interacts with WRKY40 from different plants in yeast. XopS fused to the GAL4 DNA-binding domain (BD) was expressed in combination with the WRKY40 protein from different plant species fused to the GAL4 activation domain (AD) in yeast strain Y190. Cells were grown on selective media before a LacZ filter assay was performed. Empty pGAD (AD) or pGBT9 (BD) vector served as negative control. *Nb*WRKY40, *Nicotiana benthamiana* WRKY40; *At*WRKY40, *Arabidopsis thaliana* WRKY40; *Ca*WRKY40a, *Capsicum annuum* WRKY40a. –LT, yeast growth on medium without Leu and Trp; –LTH, yeast growth on medium lacking Leu, Trp and His, indicating expression of the *HIS3* reporter gene. LacZ indicates activity of the *lacZ* reporter gene. The experiment was carried out at least two times with similar results.

WRKYs are represented by large gene families in plants and individual WRKY proteins influence diverse aspects of plant development and stress responses by acting as positive as well as negative regulators of transcription (Bakshi and Oelmüller, 2014). In order to investigate whether XopS would also bind to other members of this protein family we selected WRKY8 from *N. benthamiana*, which has been shown to be a positive regulator of defense responses (Ishihama et al., 2011) and WRKY41 from *N. tabacum* whose Arabidopsis ortholog also positively influences immunity (Higashi et al., 2008). In addition, we included *Nt*WRKY31, which we identified as an interactor of WRKY40 in a Y2H screen (Supplementary Figure S7). However, no interaction of XopS with any of the WRKYs tested could be detected in a direct Y2H assay (Supplementary Figure S7). Thus, although binding of XopS to other WRKYs not tested in this experiment cannot be excluded, there appears to be at least some specificity of XopS to interact with WRKY40 orthologs from different species.

Taken together, the Y2H experiments suggest that WRKY40 is a potential XopS target protein during a compatible interaction of *Xcv* with pepper.

### XopS interacts with WRKY40 in the plant cell nucleus

We next sought to determine whether the interactions observed in yeast would also occur *in planta*. To this end, we initially investigated the subcellular localization of WRKY40 as well as of XopS employing green fluorescent protein (GFP) – fusion proteins transiently expressed in leaves of *N. benthamiana* by infiltration with *Agrobacteria*. The fluorescence pattern was investigated using confocal laser scanning microscopy 48 hours post inoculation (hpi) of the leaves with *Agrobacteria* harboring the respective GFP-fusion construct. Expression of the fusion proteins was verified using western blotting (Figure 3A). While the XopS-GFP fluorescent signal showed a nucleo-cytoplasmic distribution, WRKY40-GFP fluorescence was confined to the nucleus of infiltrated cells (Figure 3A). Thus, subcellular localization of XopS and its potential target protein overlaps in the plant cell nucleus. To determine if XopS interacts with WRKY40 *in planta*, we performed bimolecular fluorescence complementation (BiFC) assays in *N. benthamiana*. We transiently co-expressed XopS fused with an N-terminal part of the yellow fluorescent protein derivate Venus (Venus^N173^) with WRKY40 fused with a C-terminal Venus fragment (Venus^C155^) in *N. benthamiana* leaves, which resulted in bright YFP fluorescence visualized with confocal microscopy (Figure 3B). Consistent with the nuclear localization observed for both WRKY40-GFP and XopS-GFP, the BiFC fluorescent signal was confined to the nucleus. A close-up revealed that the yellow fluorescence was not uniformly distributed inside the nucleus, but rather concentrated to discrete foci (Figure 3B). *Agrobacterium*-infiltration of XopS-Venus^N173^ together with FBPase-Venus^C155^ or FBPase-Venus^N173^ together with WRKY40-Venus^C155^ served as negative controls and did not result in fluorescence (Supplementary Figure S8). These data suggest that XopS interacts with WRKY40 inside the nucleus.

**Figure 3.**
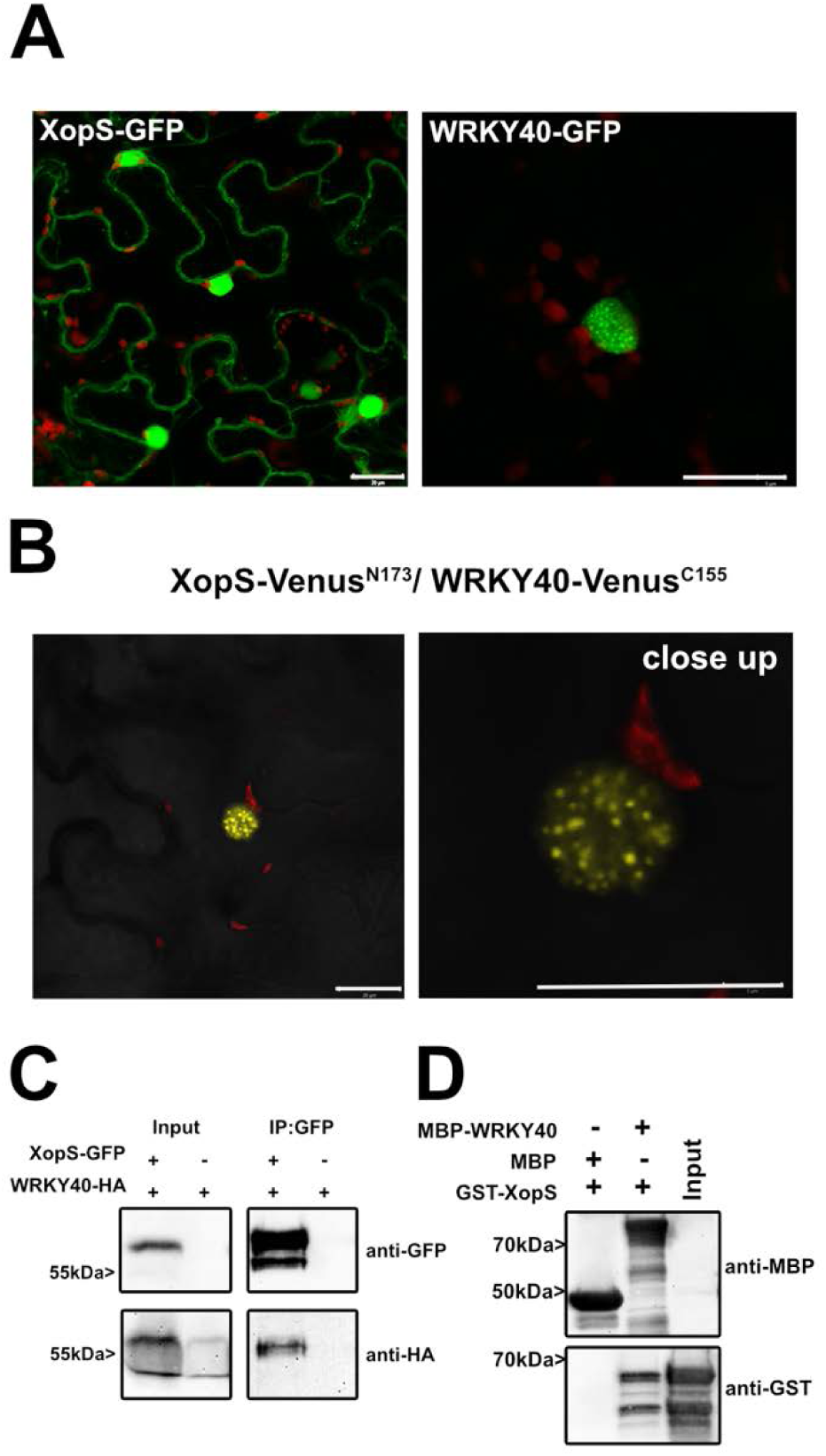
WRKY40 interacts with XopS *in planta* and *in vitro*. **(A)** Subcellular localization of XopS-GFP and *Nb*WRKY40-GFP. Green fluorescent protein (GFP) fusion proteins under control of the CaMV35S promoter were expressed transiently in leaves of *Nicotiana benthamiana* (*N. benthamiana*) using *Agrobacterium-*infiltration. The localization of transiently expressed proteins was determined with confocal laser scanning microscopy 48 hours post inoculation (hpi). Scale bars represent 20 µm. The experiment was carried out at least two times with similar results. **(B)** Visualization of protein-protein interactions *in planta* by the bimolecular fluorescence complementation assay. Yellow fluorescent protein (YFP) confocal microscopy images show *N. benthamiana* leaf epidermal cells transiently expressing XopS-Venus^N173^ in combination with *Nb*WRKY40-Venus^C155^. Pictures were taken 48 hpi. Scale bars represent 20 µm. The experiment was carried out at least two times with similar results. **(C)** Coimmunoprecipitation of XopS-GFP with *Nb*WRKY40-HA (hemagglutinin). Proteins were transiently co-expressed in leaves of *N. benthamiana* using *Agrobacterium*-infiltration. After 48 h, total proteins (Input) were subjected to immunoprecipitation (IP:GFP) with GFP-Trap^®^ beads, followed by western blot analysis using either anti-GFP or anti-HA antibodies. The experiment was carried out at least two times with similar results. **(D)** *In vitro* pull-down assay showing physical interaction of XopS with *Nb*WRKY40. Maltose-binding protein (MBP), MBP-*Nb*WRKY40 and glutathione S-transferase (GST)-XopS were expressed in *E. coli.* Pull-down was performed using amylose resin. MBP alone was used as a negative control. Proteins were detected by western blot analysis using anti-MBP or anti-GST antibodies. The experiment was carried out at least two times.

To further substantiate this finding, a GFP pull-down assay was performed. To this end, XopS-GFP was transiently co-expressed together with or without WRKY40-HA (hemagglutinin) in *N. benthamiana*. Two days after infiltration, XopS-GFP was pulled down using GFP-trap^®^ beads, and the eluates were analyzed by immunoblot with anti-GFP and anti-HA antibodies, respectively. XopS-GFP was able to pull down WRKY40-HA, verifying the interaction of both proteins *in planta* (Figure 3C).

To exclude that additional plant proteins mediate the interaction between XopS and WRKY40, an *in vitro* pull-down assay using recombinant proteins was performed. Recombinant glutathione S-transferase (GST) tagged XopS bound to a GST-affinity matrix was incubated with maltose-binding protein (MBP) tagged WRKY40. After washing, protein complex formation was analyzed by western blotting using anti-GST and anti-MBP antibodies, respectively. The western blot revealed that GST-XopS was able to pull down MBP-WRKY40, demonstrating direct physical interaction between XopS and its potential target protein without any additional eukaryotic factors (Figure 3D).

### Virus-induced gene silencing of CaWRKY40a in pepper leads to increased resistance towards Xcv

Having established that XopS directly binds to WRKY40 in plant cells, we sought to investigate the role of the transcription factor in the interaction of pepper with *Xcv*. Many WRKY TFs have been shown to respond transcriptionally to defense signaling molecules (Eulgem et al., 2000). Expression of *CaWRKY40a* was nearly undetectable in leaves from naïve pepper plants, while mRNA levels were drastically increased 10 hpi with wild type *Xcv* (Figure 4A). Induced WRKY mRNA accumulation is often extremely rapid and transient (Eulgem et al., 2000). Therefore, we monitored *CaWRKY40a* mRNA levels over a time course of 50 h after treatment of leaves with 5 mM SA. Expression peaked at 4 h after SA application to then decline rapidly by approximately 80% until 8 h post treatment (Figure 4B). Subsequently, *CaWRKY40a* mRNA levels slowly decrease further to reach almost basal levels again at the end of the time course.

**Figure 4.**
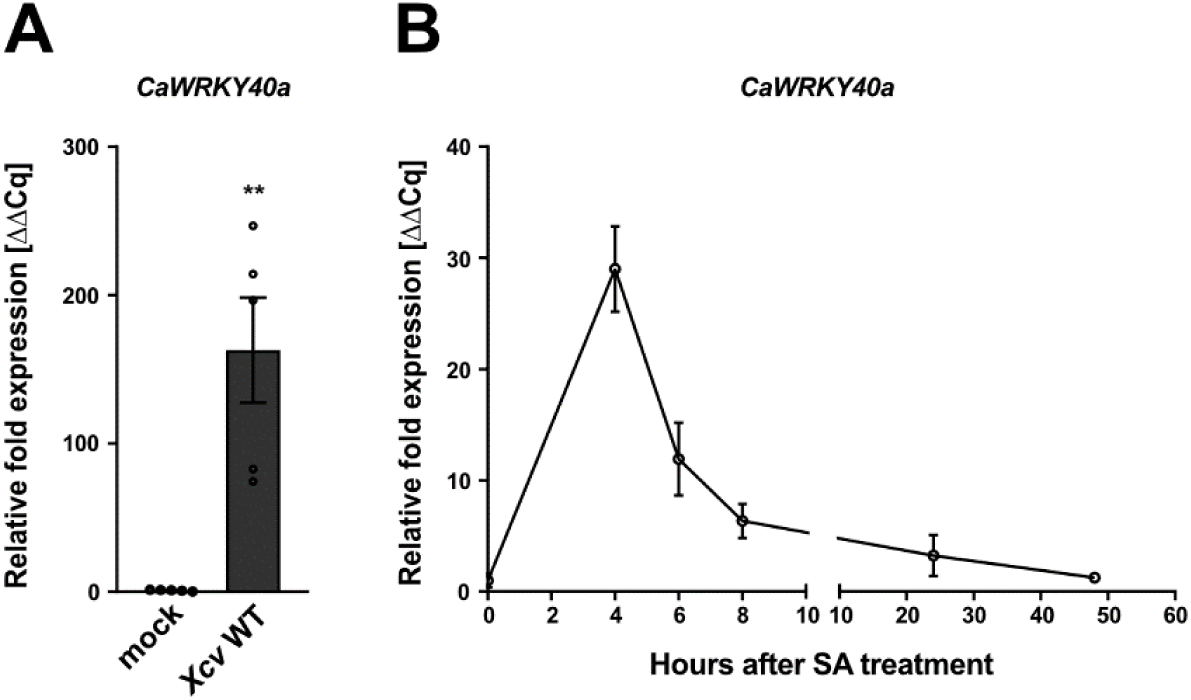
*CaWRKY40* expression is induced in response to defense signals. **(A)** Gene expression analysis of *CaWRKY40a* upon *Xcv* infection. Leaves of susceptible pepper plants were syringe-inoculated with *Xcv* wild type bacteria at OD_600_ = 0.2. Samples were taken 10 hpi, the mRNA level of *CaWRKY40a* was measured by qRT-PCR and compared to MgCl_2_ (mock) treated leaves. *Tubulin* was used as a reference gene. Each bar represents the mean of n = 5 biological replicates ± SE and asterisks (**, P < 0.01) mark significant differences according to Student’s *t*-test. The experiment was carried out three times with similar results. **(B)** Time course of *CaWRKY40a* expression in response to salicylic acid (SA). Leaves of pepper plants were sprayed with 5 mM SA (0 hours after SA treatment) and samples were taken for the relative quantification of *CaWRKY40a* expression at the indicated time points. Each data point represents the mean of n = 3 biological replicates ± SE. *Actin* was used as a reference gene. The experiment was carried out once.

To obtain direct evidence for an involvement of *CaWRKY40a* in the defense against *Xcv*, we used virus-induced gene silencing (VIGS) with Tobacco rattle virus (TRV) of *CaWRKY40a* in pepper, followed by infection with *Xcv*. For this purpose, cotyledons of pepper seedlings were infiltrated with a mixture of *Agrobacterium tumefaciens* strains of pTRV1 (Cauliflower mosaic virus; CaMV 35S-driven TRV RNA1) and pTRV2-*CaWRKY40a* (TRV RNA2 containing the target sequence), or pTRV2-*GFP* (serving as a control for infection symptoms). Four weeks after infiltration, leaves were harvested and treated with 5 mM SA for 4h to induce *CaWRKY40a* expression. Strong down regulation of *CaWRKY40a* could be confirmed by qRT-PCR in plants infiltrated with the pTRV2-*CaWRKY40a* vector as compared to the pTRV2-*GFP* control (Figure 5A). Subsequently, silenced plants were syringe-inoculated with *Xcv* and disease progression was monitored over the course of 5 days. While infected leaves of pTRV2-*GFP* plants developed strong chloroses, silencing of *CaWRKY40a* substantially reduced the appearance of visible symptoms of disease (Figure 5B). Chlorophyll content was measured as a proxy to quantify the infection phenotype. In accordance with the development of leaf chlorosis, chlorophyll content was significantly reduced in control plants as compared to *CaWRKY40* silenced plants (Figure 5C). Strikingly, bacterial multiplication in pTRV2-*CaWRKY40a* plants was significantly reduced relative to the control at 5 dpi (Figure 5D), indicating that silencing of *CaWRKY40a* in susceptible pepper plants leads to enhanced resistance toward infection with *Xcv*.

**Figure 5.**
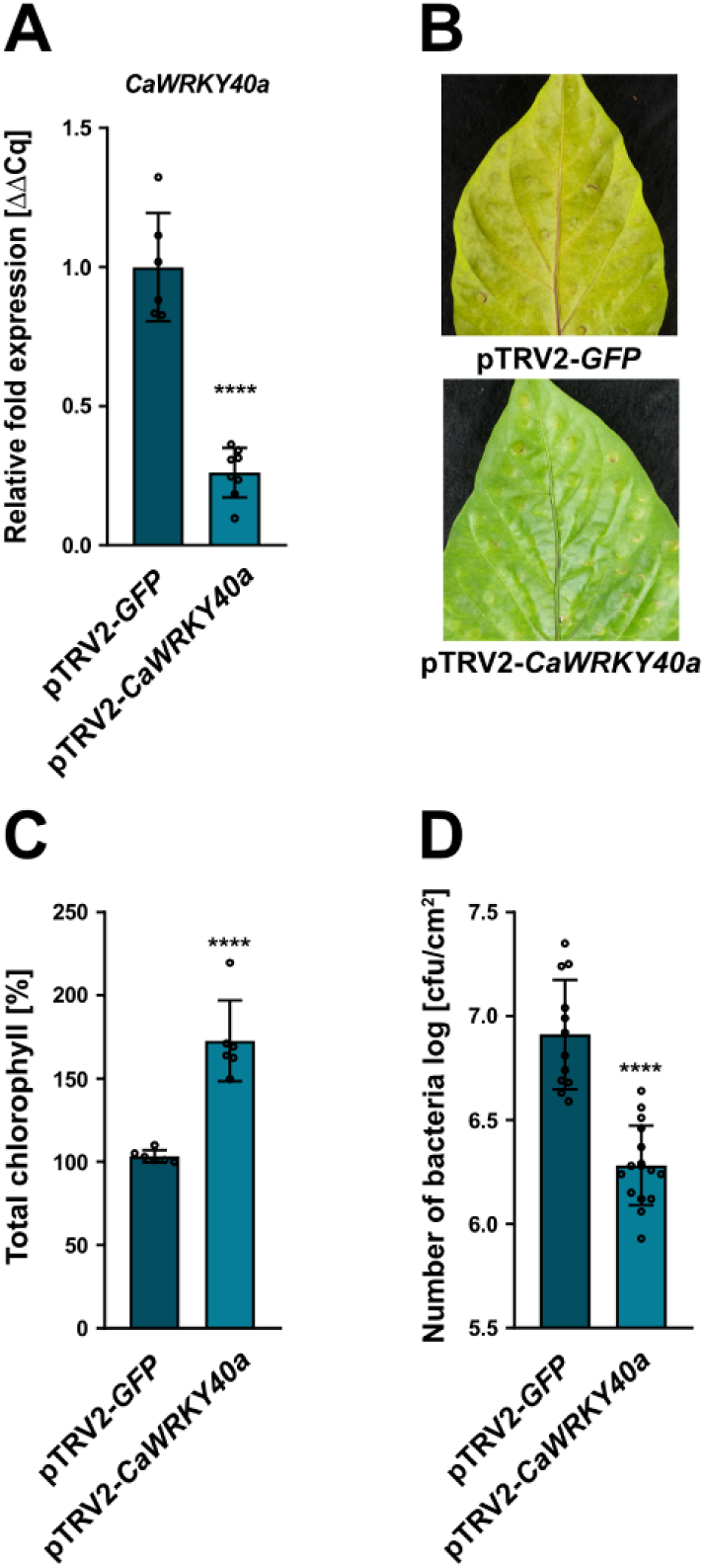
Virus-induced gene silencing of *CaWRKY40a* in pepper enhances defense against *Xcv* infection. **(A)** Verification of *CaWRKY40a* down-regulation in pTRV2-*CaWRKY40a* (*CaWRKY40a* silencing) compared to pTRV2-*GFP* (*GFP* silencing; control) in virus-induced gene silencing (VIGS) pepper plants. Three weeks after infiltrating pepper cotyledons with the silencing constructs, total RNA was isolated from excised leaves treated with 5mM SA for 4 h. The mRNA level of *CaWRKY40a* in pTRV2-*CaWRK40a* was measured by qRT-PCR and compared to *CaWRKY40a* expression in pTRV2-*GFP* control plants. *Tubulin* was used as a reference gene. Bars represent the mean of at least n = 6 biological replicates ± SD (n = 6 for pTRV2-*GFP* and n = 8 for pTRV2-*CaWRKY40a*). Asterisks (****, P < 0.0001) mark significant differences according to Student’s *t*-test. The experiment was carried out three times with similar results. **(B)** Disease symptom development in leaves of pTRV2-*CaWRKY40a* pepper plants compared to pTRV2-*GFP* control plants, syringe-inoculated with wild type *Xcv* bacteria at an OD_600_ = 0.0001. Pictures were taken at 5 dpi. A similar phenotype was observed every time plants were syringe-inoculated with *Xcv*. **(C)** Chlorophyll levels in leaves of pTRV2-*CaWRKY40a* pepper plants syringe infected with wild type *Xcv* relative to infected pTRV2-*GFP* plants. The percentage of total chlorophyll was determined when compared to the pTRV2-*GFP* control (set to 100%). The measurement was performed 5 dpi. Each bar represents the mean of n = 6 biological replicates ± SD and asterisks (****, P < 0.0001) mark significant differences according to Student’s *t*-test. The experiment was carried out twice with similar results. **(D)** Bacterial growth of wild type *Xcv* in leaves of pTRV2-*CaWRKY40a* compared to pTRV2-*GFP* control plants. Leaves were syringe-inoculated with a bacterial density of OD_600_ = 0.0001 and colony forming units in infected tissue were quantified at 5 dpi. Bars represent the mean of at least n = 6 biological replicates (and 2 technical replicates per biological replicate) ± SD (n = 6 for pTRV2-*GFP* and n = 8 for pTRV2-*CaWRKY40a*). Asterisks (****, P < 0.0001) mark significant differences according to Student’s *t*-test. The experiment was carried out three times with similar results.

### The ability of XopS to interfere with stomatal immunity requires WRKY40

To investigate whether the ability of XopS to prevent stomatal closure in response to a MAMP-stimulus is dependent on WRKY40, we used VIGS to down-regulate *NbWRKY40* expression in *N. benthamiana*. Three weeks after co-infiltration of leaflets with pTRV1 and pTRV2-*NbWRKY40* or pTRV2 (empty vector; EV control), excised leaves were treated with 5 mM SA to induce WRKY40 expression and the silencing effect was quantified using qRT-PCR. As shown in Figure 6A, VIGS resulted in a reduction of *WRKY40* gene expression of app. 90% as compared to the pTRV2 control. Subsequently, XopS-GFP or GFP alone were expressed in leaves of *WRKY40* as well as EV silenced plants using *Agrobacterium*-infiltration (Supplementary Figure S9). 24 hpi leaf discs of XopS-GFP expressing plants along with GFP controls were monitored for the ability to close their stomata in response to flg22. Measurement of stomatal aperture 2 h post treatment revealed that stomata significantly closed in EV control plants expressing GFP as compared to those infiltrated with XopS-GFP. This confirms the ability of XopS to attenuate stomatal closure in response to a MAMP stimulus (Figure 6B). pTRV2*-NbWRKY40* plants showed comparable stomatal closure in response to flg22 exposure as control plants. In turn, expression of XopS-GFP had no significant effect on stomatal aperture in plants with reduced *WRKY40* expression (Figure 6B). These data clearly demonstrate a direct relationship of WRKY40 and the ability of XopS to interfere with stomatal immunity.

**Figure 6.**
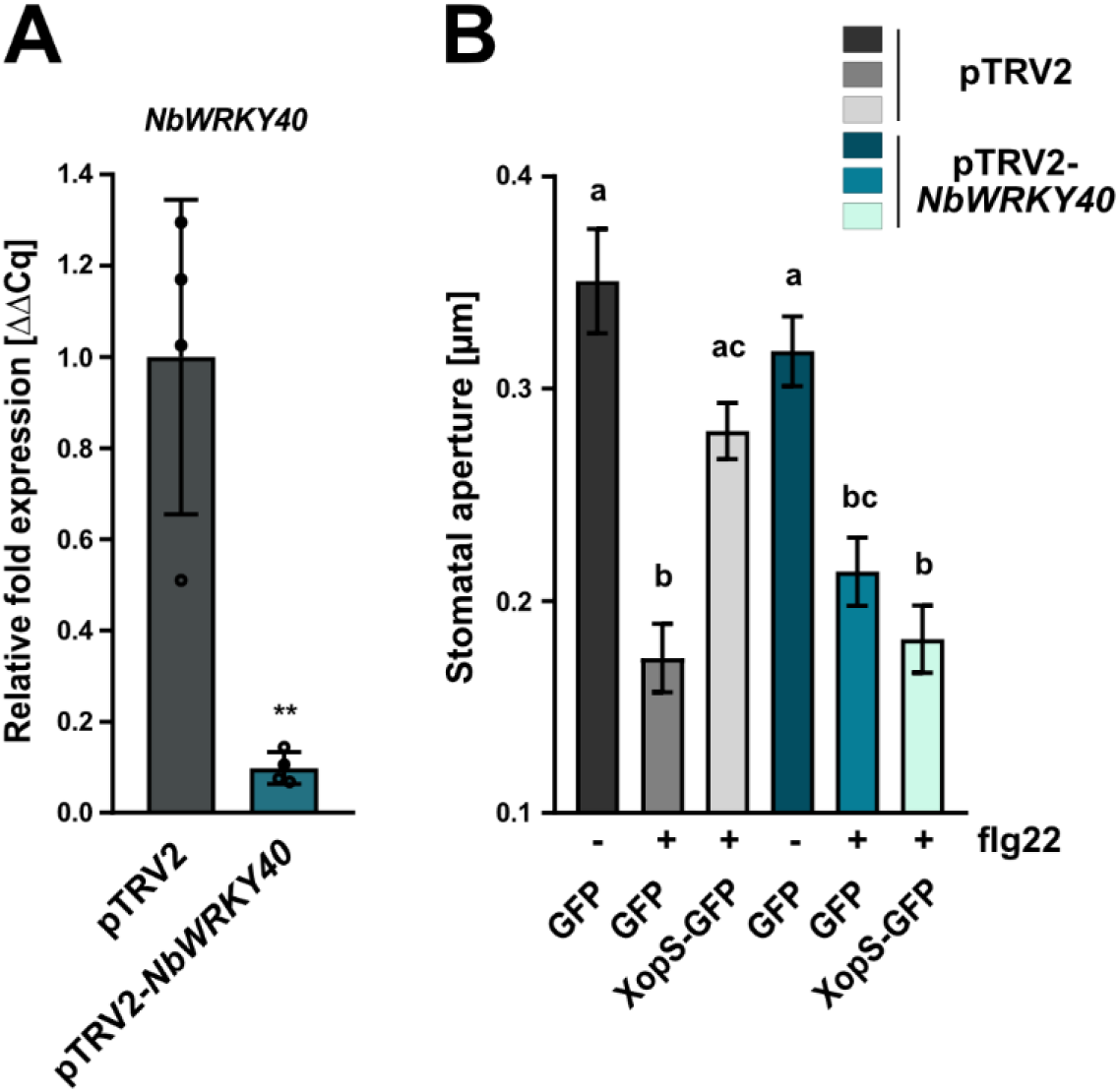
Virus-induced gene silencing of *WRKY40* in *N. benthamiana* affects stomatal closure in response to a MAMP stimulus. **(A)** Verification of *NbWRKY40* down-regulation in pTRV2-*NbWRKY40* (*NbWRKY40* silencing) compared to pTRV2 (empty vector, silencing control) in VIGS *N. benthamiana* plants. Two weeks after infiltrating *N. benthamiana* plants with the silencing constructs, total RNA was isolated from excised leaves treated with 5mM SA for 4 h. The mRNA level of *NbWRKY40* in pTRV2-*NbWRK40* was measured by qRT-PCR and compared to *NbWRKY40* expression in pTRV2 control plants. *Actin* was used as a reference gene. Bars represent the mean of n = 4 biological replicates ± SD and asterisks (**, P < 0.01) mark significant differences according to Student’s *t*-test. **(B)** Stomatal aperture measurement in pTRV2-*NbWRKY40* and pTRV2 *N. benthamiana* plants transiently expressing either GFP alone or XopS-GFP. Leaf discs were floated on water (control) or on water supplemented with 25 µM flg22 for two hours prior to the measurement of stomatal aperture under a microscope. Approximately 100 apertures from n = 4 independent plants were measured per individual treatment and are represented as width/length ratio. Bars represent the mean ± SE. Letters over bars represent statistical significance determined by one-way ANOVA (P < 0.05). The experiment was carried out twice with similar results.

### WRKY40 negatively regulates defense gene expression

The enhanced resistance of pTRV2-*CaWRKY40a* plants to infection by *Xcv* suggests that similar to the scenario in Arabidopsis, the TF is involved in negative regulation of defense gene expression. To investigate if XopS interacting WRKY40s are capable of transactivation, we performed a yeast transactivation assay (Ye et al., 2004). To this end the full length ORFs of *NbWRKY40* and *CaWRKY40a* were cloned into the pGBT9 vector to create fusion proteins with the GAL4 DNA-binding domain (BD). *NbWRKY8*, a known positive regulator of gene expression (Ishihama et al., 2011) served as a control.

The vectors were separately transformed into the Y190 yeast reporter strain and transformants were assayed for growth on histidine lacking medium and for *LacZ*-activity. While BD-*Nb*WRKY8 was able to induce strong reporter gene expression, BD-*Nb*WRKY40 and BD-*Ca*WRKY40a did not display transactivation activity in yeast (Figure 7A). In order to confirm the repressor activity of WRKY40 *in planta*, we fused a GUS reporter gene to a nucleotide sequence containing 4 W-boxes in front of a minimal CaMV35S promoter (Figure 7B). Co-expression of the *W-box::GUS* construct with CaMV35S driven *NbWRKY8* resulted in significant induction of GUS activity relative to the empty vector (EV) control (Figure 7C), which is in line with a role of WRKY8 as a positive regulator of gene expression (Ishihama et al., 2011). In contrast, co-expression with CaMV35S driven WRKY40 significantly repressed GUS activity below the level of the EV control, suggesting that WRKY40 is able to suppress basal GUS expression mediated by the minimal CaMV35S promoter (Figure 7C). Expression of both WRKY proteins tested was verified by western blotting (Supplementary Figure S10). Thus, WRKY40 likely functions as negative regulator of gene expression.

**Figure 7.**
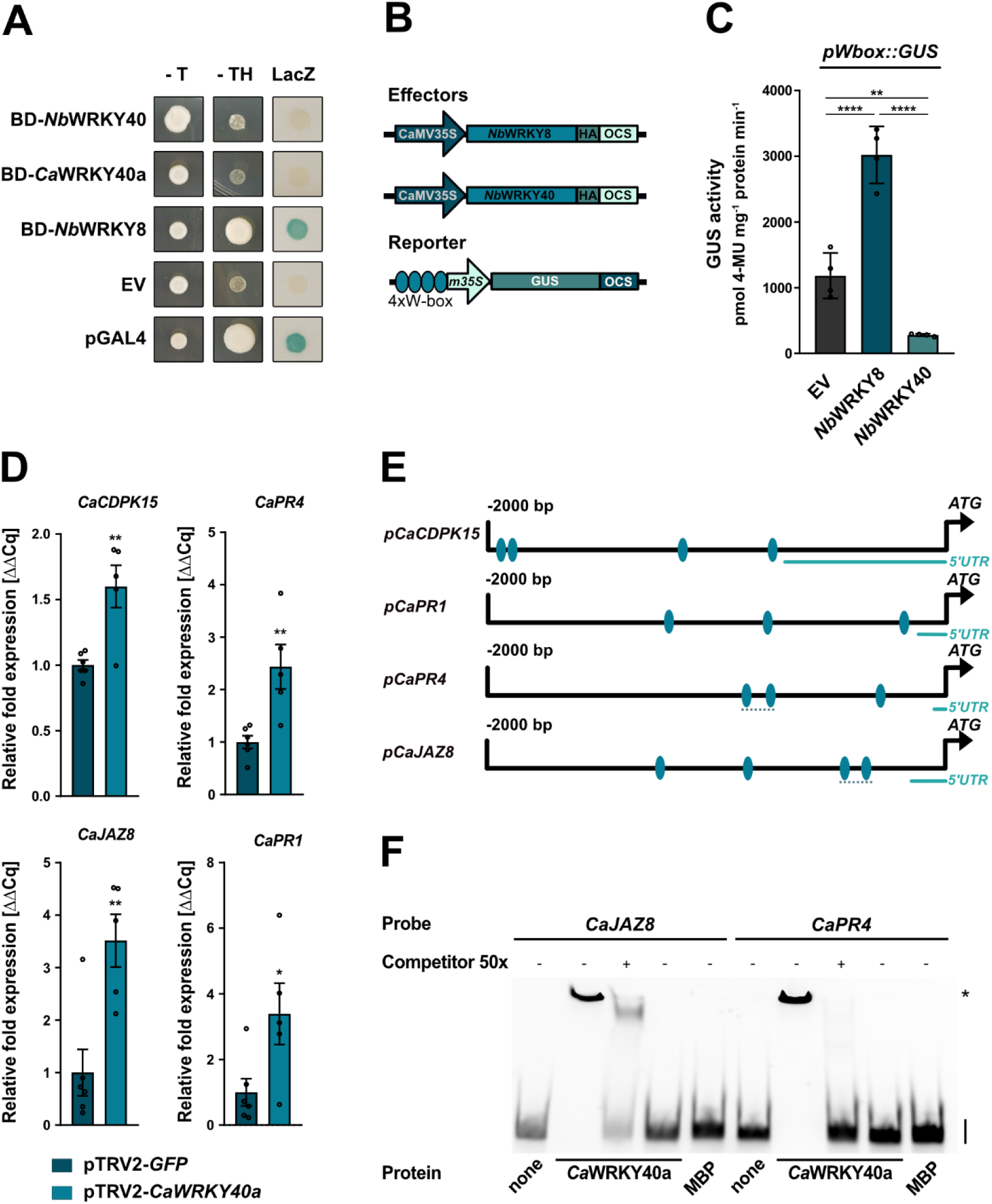
WRKY40 is a negative regulator of defense gene expression. **(A)** Yeast transactivation assay. WRKY proteins were expressed as GAL4 DNA-BD fusion proteins in yeast strain Y190. *HIS3* and *LacZ* reporter gene activity was monitored 3 days after spotting transformed cells on selective medium. The empty pGBT9 vector (EV) served as negative control and pGAL4 vector (pGAL4) containing full length GAL4 was used as positive control. – T, medium lacking Trp; - TH, medium lacking Trp and His, indicating expression of the *HIS3* reporter gene. LacZ indicates activity of the *lacZ* reporter gene. *Nb*WRKY40, *Nicotiana benthamiana* WRKY40; *Ca*WRKY40a, *Capsicum annuum* WRKY40a; *Nb*WRKY8, *N. benthamiana* WRKY8. The experiment was carried out three times with similar results. **(B)** Schematic representation of the effector and reporter constructs used in the transient expression assay of W-box driven promoter activity. Effector constructs contained either *Nb*WRKY8 or *Nb*WRKY40 under control of the constitutive CaMV35S promoter. A C-terminal HA-tag enabled immune detection of the protein. The *β-Glucuronidase* (*GUS*) reporter gene was submitted to the control of minimal CaMV35S promoter (*m35S*) preceded by an oligonucleotide containing 4 W-boxes (TTGAC(C/T). OCS = terminator of the octopine synthase. **(C)** W-box driven GUS reporter gene activity in *N. benthamiana*. The GUS reporter gene was submitted to the control of a chimeric promoter consisting of a 4 x W-box element fused to the minimal CaMV35S promoter. The reporter construct was transiently co-expressed with empty vector HA (EV), *Nb*WRKY8-HA or *Nb*WRKY40-HA as effector constructs. GUS activity was determined 48 h post *Agrobacterium*-infiltration and is represented as pmol 4-Methylumbelliferone mg protein^-1^ min^-1^. Bars represent the mean of n = 4 biological replicates ± SD and asterisks (**, P < 0.01; ****, P < 0.0001) mark significant differences according to one-way ANOVA. The experiment was carried out three times with similar results. **(D)** Defense gene expression in pTRV2-*CaWRKY40a* pepper plants upon infection with *Xcv*. Leaves of pTRV2-*CaWRKY40a* and pTRV2-*GFP* control pepper plants were syringe-inoculated with *Xcv* wild type bacteria at OD_600_ = 0.2 and samples were taken 10 hpi. The mRNA levels of four indicated potential *Ca*WRKY40a target genes were measured by qRT-PCR in pTRV2-*CaWRKY40a* plants and were compared to their mRNA levels in pTRV2 control plants. *Ubiquitin-conjugating protein* (*UBI*-3) was used as a reference gene. Bars represent the mean of at least n = 5 biological replicates ± SE (n = 6 for pTRV2-*GFP* and n = 5 for pTRV2-*CaWRKY40a*). Asterisks (*, P < 0.05; **, P < 0.01) mark significant differences according to Student’s *t* test. The experiment was carried out twice with similar results. **(E)** Schematic representation of the 2 kb promoter regions upstream of the translational start site of indicated candidate *Ca*WRKY40a target genes. Ellipsis indicate the position of predicted W-boxes. Dashed lines represent position of the probes for electrophoretic mobility shift assay (EMSA). **(F)** EMSA shows binding of *Ca*WRKY40a to W-boxes contained in the *CaJAZ8* and *CaPR4* promoter regions. MBP-*Ca*WRKY40a was produced in *E. coli* and incubated with Cy5-labelled 150 bp DNA fragments derived from the *CaJAZ8* and *CaPR4* promoter regions. Each DNA fragment contained two predicted W-boxes. Protein-DNA complexes were separated from unbound probe on a 5% TBE gel. Unlabeled DNA fragment was used as a competitor with a 50 fold (50x) excess over the amount of used probe. MBP protein alone was included as additional negative control. At the right-hand side of the gel, specific retarded protein-DNA complexes are marked by an asterisk, whereas free running probes are designated by a black bar. The experiment was carried out twice with similar results.

We next analyzed pathogen-induced defenses gene expression in pTRV2-*CaWRKY40a* plants compared to control plants. Since target genes of *Ca*WRKY40a in pepper are not known, we chose genes which were described earlier to be under the regulation of WRKY TFs, such as *CaCDPK15* (Shen et al., 2016) and *CaPR4* (Huh et al., 2015), or were considered as *bona fide* WRKY target genes based on their regulation in Arabidopsis such as *JAZ8* and *PR1* (Pandey et al., 2010; Birkenbihl et al., 2017). As a prerequisite for WRKY binding, all selected genes were confirmed to contain W-boxes ((T)TGAC(C/T)) within the 2 kb upstream region relative to their transcriptional start site (Figure 7E). When measured 10 h after syringe-infection with wild type *Xcv* bacteria, the expression of all genes tested was significantly higher in pTRV2-*CaWRKY40a* plants as compared to the pTRV2-GFP silencing control (Figure 7D), suggesting that silencing of *CaWRK40a* leads to a faster and/or stronger defense response on the transcriptional level correlating with enhanced *Xcv* resistance. Effective silencing of *CaWRKY40a* in the plants under scrutiny was verified by qRT-PCR (Supplementary Figure S11).

We next sought to investigate whether increased expression level of the genes was due to the loss of *Ca*WRKY40a binding to their promoter regions. To this end, direct binding of recombinant MBP-*Ca*WRKY40a to an app. 150 bp *CaJAZ8* or *CaPR15* promoter fragment comprising two predicted W-boxes (Figure 7E) was analyzed using an electrophoretic mobility shift assay (EMSA).

The results shown in Figure 7F indicate that the MBP-*Ca*WRKY40a protein was able to bind to CY5-labelled promoter fragments containing the W-boxes resulting in a shift of mobility. Excess unlabeled probe (the same DNA fragment without CY5) efficiently competed for binding to the protein, indicating specificity of binding. No binding signal was detected in reactions without added protein (lane 1 and 6) or with MBP alone (lane 5 and 10). Thus, *CaJAZ8* and *CaPR4* are likely subject to direct regulation by *Ca*WRKY40a in pepper plants.

### XopS interferes with proteasomal turnover of WRKY40 *in planta*

Having established the interaction between XopS and WRKY40 and the role of the TF as a negative regulator of defense gene expression, we next sought to determine how XopS might affect the function of WRKY40 to support bacterial virulence. Some WRKY TFs have been shown to undergo proteasomal turnover to regulate their activity (Miao and Zentgraf, 2010; Matsushita et al., 2013; Ye et al., 2018; Liu et al., 2021). Based on the initial observation that WRKY40 protein levels in *Agrobacterium*-mediated transient expression assays were comparatively low, we set out to determine whether WRKY40 undergoes proteasomal degradation *in planta*.

To this end, an HA-tagged version of WRKY40 was transiently expressed in leaves of *N. benthamiana*. After two days the same leaves were infiltrated with the well-characterized proteasome inhibitor MG132 and samples were taken over a time course of 6h. As shown in Figure 8A, at 0 h WRKY40-HA protein was only detectable after long exposure of the membrane while the signal was barely visible upon shorter exposure. However, the WRKY40-HA signal increased gradually over time becoming easily visible also upon short exposure 6 h after treatment with MG132, indicating accumulation of WRKY40-HA when the proteasome is inhibited. The long exposure revealed the presence of higher molecular weight WRKY40-HA species at later time points of MG132 treatment reminiscent of a poly-ubiquitination of the protein (Figure 8A). From these data we conclude that WRKY40 undergoes proteasomal degradation when transiently expressed in *N. benthamiana*.

**Figure 8.**
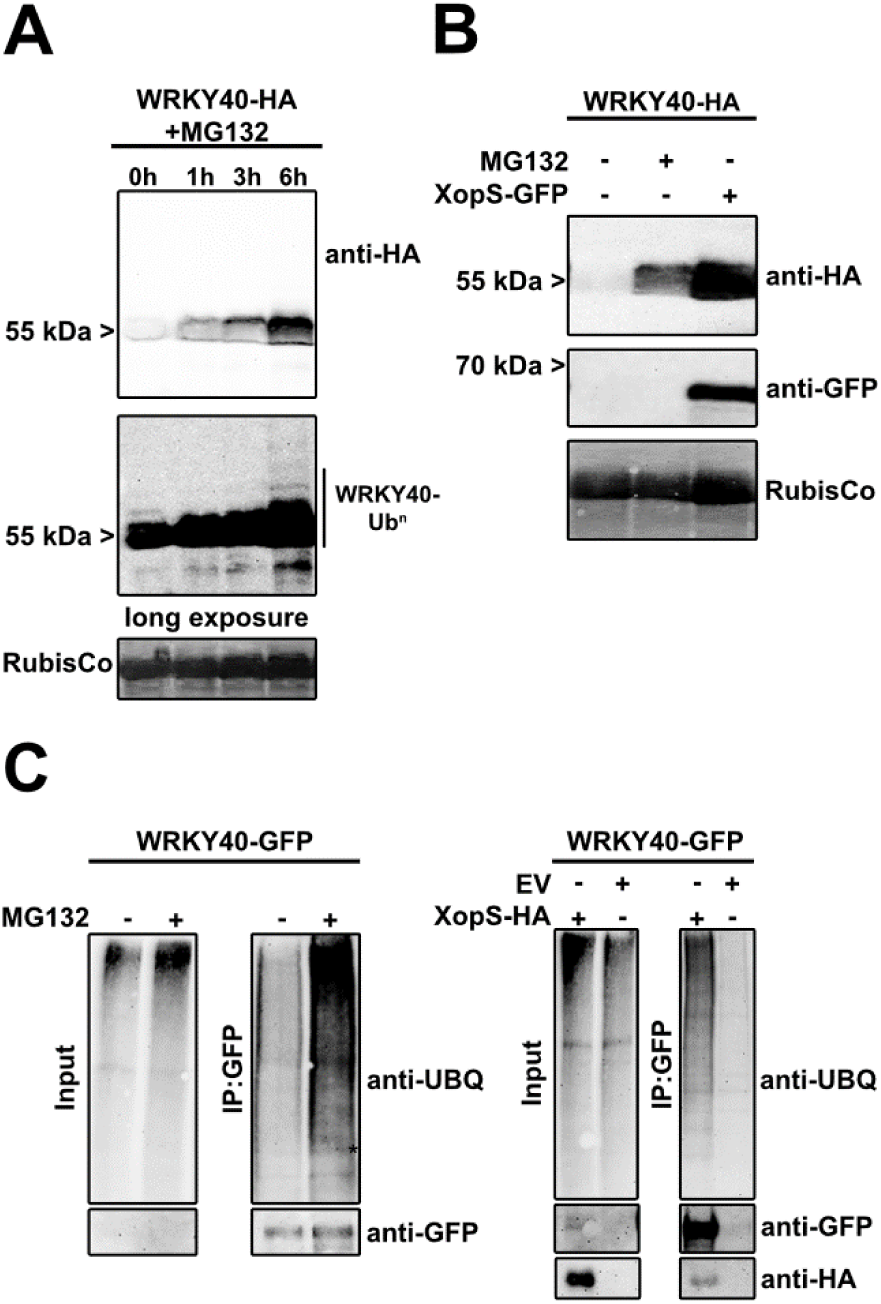
XopS protects WRKY40 from degradation. **(A)** Inhibition of the 26S proteasome results in the accumulation of WRKY40. *Nb*WRKY40-HA was transiently expressed in leaves of *N. benthamiana* using *Agrobacterium*-infiltration. 48 hpi leaves were treated with 200 µM MG132 and samples were taken at the time points indicated. *Nb*WRKY40-HA protein levels were monitored over time using an anti-HA antibody (upper panel: short exposure; lower panel: long exposure). Amido black staining of RubisCo served as a loading control. The experiment was carried out at least two times with similar results. **(B)** Co-expression with XopS-GFP results in increased *Nb*WRKY40-HA protein levels. *Nb*WRKY40-HA was transiently expressed either alone or along with XopS-GFP or was treated with 200 µM MG132 for 6 h where indicated. Proteins were detected using either anti-HA or anti-GFP antibodies 48 hpi. Amido black staining of RubisCo served as loading control. The experiment was carried out at least two times with similar results. **(C)** Accumulation of ubiquitinated *Nb*WRKY40-GFP in the presence of XopS. *Nb*WRKY40-GFP was transiently expressed and the protein was stabilized either by treatment with 200 µM MG132 or by co-expression with XopS-HA (Input). Subsequently, the protein was purified from total plant extracts using GFP-trap^®^ (IP:GFP). Proteins were detected by western blotting with either anti-Ubiquitin, anti-GFP or anti-HA antibodies. The experiment was carried out at least two times with similar results.

To explore whether XopS affects WRKY40 protein accumulation, WRKY40 levels were monitored by western blotting in leaves of *N. benthamiana* either expressing WRKY40 alone or in combination with XopS. As previously observed, WRKY40-HA was only readily detectable when the proteasome was inhibited by MG132, indicative for its proteasomal turnover (Figure 8B). However, when co-expressed with XopS-GFP, WRKY40 accumulated to high levels suggesting that the effector renders WRKY40 resistant to proteasomal degradation (Figure 8B). No effect on overall proteasome activity could be detected in leaves transiently expressing XopS (Supplementary Figure S12), indicating that the effector has no general inhibitory effect on the proteasome as previously reported for other T3Es (Üstün et al., 2013a; Üstün et al., 2014; Üstün et al., 2016). Furthermore, a recent study also suggest that XopS does not influence autophagy, another protein degradation pathway (Leong et al., 2021). Most proteasome substrates are marked for degradation by ubiquitin conjugation (Vierstra, 2009). These conjugates can be reversed or modified by de-ubiquitinating enzymes (DUBs) which could interfere with the proteasomal degradation of target proteins. In order to investigate whether XopS affects the overall ubiquitination status of WRKY40 we expressed a WRKY40-GFP fusion protein transiently in leaves of *N. benthamiana* and inhibited its proteasomal degradation either by MG132 treatment or by co-expression with XopS. Subsequently the protein was immunopurified using a GFP-trap^®^ and monitored for ubiquitination using an ubiquitin-specific antibody. As shown in Figure 8C, inhibition of WRKY40 proteasomal degradation either by MG132 or by XopS allowed purification of WRKY40-GFP in its ubiquitinated form. This suggests that XopS does not stabilize WRKY40 by preventing its ubiquitination through e.g. de-ubiquitination. However, the assay does not allow to draw conclusions about WRKY40 ubiquitination levels or ubiquitin chain topology.

Given the role of WRKY40 as a negative regulator of defense gene expression, we hypothesized that stabilization of WRKY40 through interaction of XopS would enhance the repressor function of the TF. To test this, we fused the 2 kb upstream region of the *CaPR4* gene (*pCaPR4*) to a GUS reporter gene and co-transfected the construct along with CaMV35S driven *NbWRKY40* or *NbWRKY8* either in the absence or presence of XopS in leaves of *N. benthamiana* (Figure 9A). The *pCaPR4* fragment mediated readily detectable GUS expression in the presence of the EV control (Figure 9B). Expression of the effector constructs was verified by western blotting (Supplementary Figure S13). XopS alone already led to a significant reduction of GUS activity, likely through stabilization of endogenous WRKY40 protein. WRKY8 alone had no significant effect on *pCaPR4* driven GUS expression; however, in the presence of the positive regulator of gene expression WRKY8, XopS was not able to cause a reduction in reporter gene activity. Transient expression of WRKY40 led to a strong repression of *pCaPR4* driven GUS activity which was slightly but not significantly enhanced in the presence of XopS (Figure 9B).

**Figure 9.**
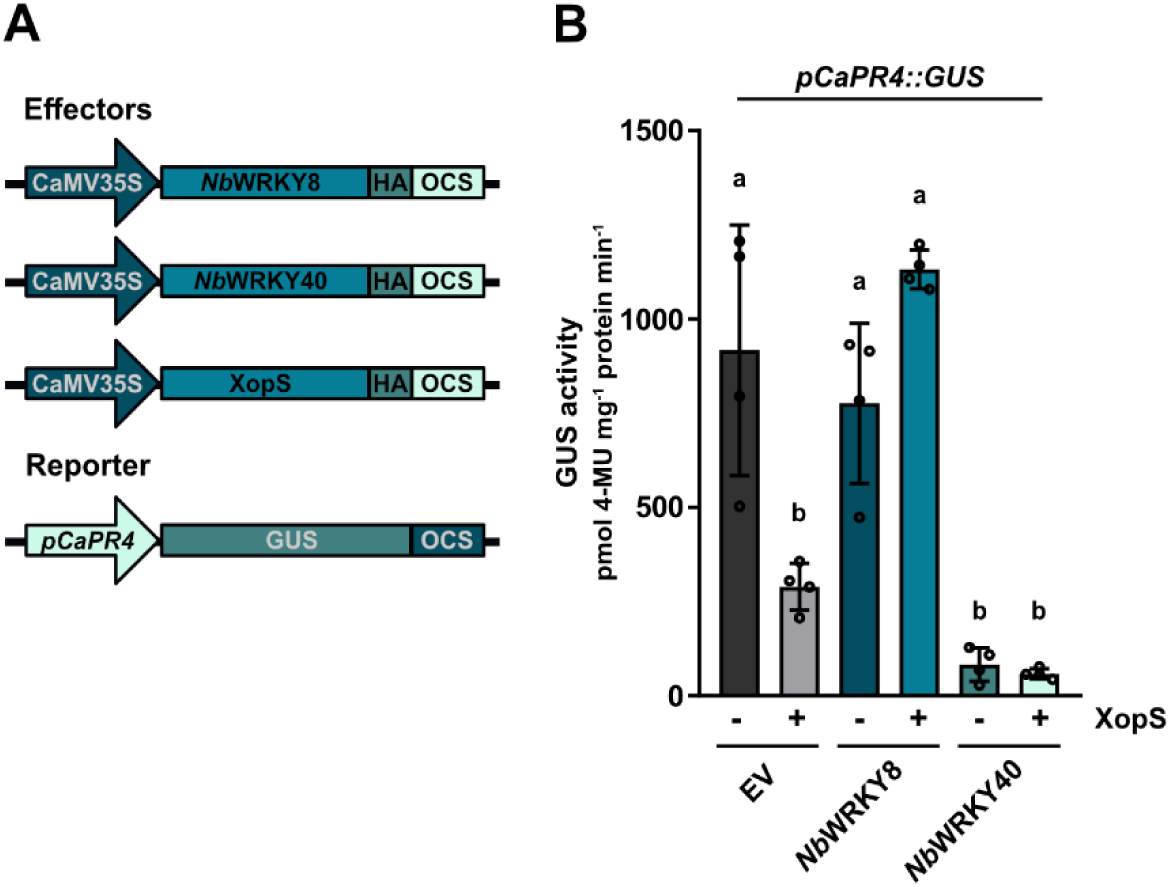
XopS and WRKY40 repress expression from the *CaPR4* promoter. **(A)** Schematic diagrams of the effector and reporter constructs. The effector plasmids contain the WRKY TFs or XopS fused to the constitutive CaMV35S promoter and carry a C-terminal HA-tag for immunodetection of the protein. The GUS reporter construct contains the 2 kb region upstream of the predicted translational start site of the *CaPR4* gene. **(B)** Transactivation of the *pCaPR4*::GUS reporter by the TFs *Nb*WRKY8 or *Nb*WRKY40 either in presence or absence of XopS. EV = empty vector control. Samples were taken 48 hours after *Agrobacterium*-infiltration and GUS activity is expressed in pmole 4-Methylumbelliferone mg^-1^ protein min^-1^. Bars represent the mean of n = 4 biological replicates ± SD. Letters over bars represent statistical significance determined by one-way ANOVA (P < 0.05).

## Discussion

Our data show that an *Xcv*Δ*xopS* strain achieves significantly lower population densities than wild type *Xcv* when bacteria were dip-inoculated onto the leaf surface. This is in contrast to what has been described for pressure infiltration of leaf tissue where an *Xcv* strain lacking XopS was not affected in bacterial multiplication relative to the wild type (Schulze et al., 2012). It strongly resembles the infection of Arabidopsis with *Pst cor^-^* mutant bacteria (Mittal and Davis, 1995) and suggests that similar to COR XopS interferes with an early defense response which is bypassed by syringe-inoculation of bacteria directly into the apoplast. Indeed, incubation of pepper leaves with *Xcv* lacking XopS led to considerable stomatal closure while the wild type strain was able to prevent closing of the stomata. Similar results were obtained upon flg22 treatment of either *N. benthamiana* or Arabidopsis leaves ectopically expressing XopS. This strongly suggests that XopS is either able to prevent stomatal closure or is leading to reopening of stomata after their initial closure to perpetuate bacterial tissue entry. It is not known whether the T3SS of *Xcv* is functional prior to tissue entry in order to deliver T3Es directly into guard cells or whether it influences stomatal immunity from non-stomatal cells. Zhang et al. (2009) used an *Xcv hrp-gfp* reporter strain to monitor formation of the T3SS during the epiphytic growth phase and their data suggest that moderate *hrp* gene expression occurred in cells on the leaf surface primarily in the immediate vicinity of stomata. Thus, a direct translocation of T3Es, including XopS, into guard cells by *Xcv* before tissue entry is a possible scenario.

Although deletion of XopS does not affect bacterial multiplication within the apoplast of infected pepper plants (Schulze et al., 2012), the effector is required for full symptom development and interferes with apoplastic immune responses. Like COR, XopS is required to cause chlorosis development and leads to the reduction of SA levels resulting in decreased expression of SA-dependent genes. Syringe-inoculation of a non-pathogenic *Pst* strain into the leaves of Arabidopsis lines expressing XopS demonstrates that the effector supports bacterial growth and thus interferes with PTI during apoplastic defense. Moreover, XopS expression in Arabidopsis leads to the development of leaf chlorosis, the induction of early JA-responsive genes and elevated JA levels. This phenotype is reminiscent of Arabidopsis lines expressing the T3E HopX1 from *Pseudomonas syringae* pv. *tabaci* (*Pta*) 11528, which is a protease that degrades JAZ proteins, a key family of JA-repressors (Gimenez-Ibanez et al., 2014). *Pta* does not produce COR and thus it uses HopX1 as an alternative means to promote activation of JA-induced defenses and susceptibility in Arabidopsis. Based on the congruence of XopS related phenotypes with those produced by COR and HopX1, respectively, we propose that *Xcv* uses XopS to activate JA signaling and to repress SA-related defense responses in order to promote disease susceptibility.

We show by several independent methods that XopS interacts with WRKY40 TFs inside the plant cell nucleus, suggestive for a role of XopS in modulating host gene expression. In general, WRKY factors fulfill essential regulatory functions in PTI, ETI and systemic acquired resistance (SAR) responses, acting as activators or repressors of gene expression (Rushton et al., 2010). Growing evidence suggests that WRKY TFs form interaction networks in which individual WRKY proteins make additive, cooperative or antagonistic contributions to the overall transcriptional activity of a given TF complex (Chi et al., 2013).

In Arabidopsis, WRKY40 functions redundantly with WRKY18 and WRKY60 as negative regulator of resistance against the hemibiotrophic bacterial pathogen *Pst* DC3000 (Xu et al., 2006). While neither the *wrky40* nor the *wrky18* single mutant displayed increased resistance against *Pst*, a *wrky40 wrky18* double mutant showed reduced bacterial multiplication, which was even further diminished in a *wrky40 wrky18 wrky60* triple mutant line (Xu et al., 2006). In turn, transgenic Arabidopsis lines co-expressing *WRKY40* and *WRKY18* under control of the CaMV35S promoter supported 10 times more bacterial growth than wild type plants. Enhanced bacterial resistance in *wrky* knock-out lines was accompanied by elevated expression levels of *PR1* upon infection. Similar to the *wrky40 wrky18* knock-out in Arabidopsis, VIGS of *CaWRKY40a* in pepper plants led to a more than 10-fold reduction of bacterial multiplication upon *Xcv* infection suggesting enhanced apoplastic defense against this hemibiotrophic pathogen. Reduced bacterial growth was correlated with enhanced expression of defense related genes, further corroborating a role of *Ca*WRKY40a as a negative regulator of gene expression. The *in vivo* targets of *Ca*WRKY40a currently remain unknown. Based on previously reported positive regulation by other WRKY TFs and the *in vitro* binding of *Ca*WRKY40a to W-boxes within their promoter region, *CaJAZ8* and *CaPR4* can be considered as *bona fide Ca*WRKY40a target genes. However, indirect effects by other WRKY proteins or TFs from other families on gene expression cannot be ruled out at this point. A recent genome wide Chip-seq analysis identified more than 1.400 *in vivo* target genes for WRKY40 in Arabidopsis with a clear preference for binding gene loci involved in early PTI perception and signaling (Birkenbihl et al., 2017). Thus, it can be expected that silencing of *CaWRKY40a* in pepper has far reaching consequences on defense gene expression resulting in enhanced immunity towards *Xcv*.

The fact that *CaWRKY40a* silencing alone has a substantial effect on *Xcv* resistance and defense gene expression suggests a more central role of the TF as a negative regulator of defense responses in pepper than its ortholog has in Arabidopsis. Whether and how *Ca*WRKY40a is functionally connected to other WRKY proteins is not known but given the interaction *Nb*WRKY40 with *Nt*WRKY31 in yeast it is highly likely that in pepper the TF is also involved in a higher order WRKY network. Similar to silencing of *CaWRKY40a*, VIGS of *CaWRKY1* in pepper led to a reduction in bacterial growth upon infection with *Xanthomonas axonopodis* pv. *vesicatoria* race 1, suggesting that both TFs act as negative regulators of immunity in a non-redundant but overlapping manner (Oh et al., 2008).

Several lines of evidence suggest that *Ca*WRKY40a is negatively regulating defense gene expression, including the lack of transactivation activity in yeast and the ability to suppress the expression of a GUS reporter gene driven by W-box containing promoter sequence. Thus, its induction by the defense hormone SA and during infection with virulent *Xcv* bacteria appears counterintuitive. A similar expression pattern has been observed for Arabidopsis *WRKY40* and it has been proposed that this mode of regulation could ensure a dynamic function by which some early PTI responses are rapidly activated but subsequently dampened to avoid unnecessary prolonged resource allocation to immunity (Pandey et al., 2010).

*CaWRKY40a* silenced plants display substantially reduced disease symptoms upon infection with *Xcv* compared to wild type plants, which in turn developed severe chloroses correlated with losses in chlorophyll content.

The development of chloroses is largely dependent on the translocation of XopS and the phenotypic similarities between *XcvΔxopS* infected wild type pepper plants and the *Xcv* infected *CaWRKY40a* silenced plants strongly suggests that XopS requires *Ca*WRKY40a to exert its virulence function. XopS is not able to prevent stomatal closing in *WRKY40* silenced *N. benthamiana* in response to a MAMP stimulus, reinforcing a direct link between XopS’ virulence functions and its interaction with WRKY40. The development of chloroses accompanied by a loss in chlorophyll content is considered a hallmark response of the JA pathway (Creelman and Mullet, 1995). In Arabidopsis, WRKY40 targets the promoter region of several JA associated genes, including *JAZ8* (Pandey et al., 2010; Birkenbihl et al., 2017). Although *wrky40 wrky18* and *wrky40 wrky18 wrky60* mutants display increased resistance against biotrophic and hemi-biotrophic pathogens (Xu et al., 2006; Pandey et al., 2010), they show enhanced susceptibility to the necrotrophic pathogen *Botrytis cinerea* correlated with a reduced JA response (Xu et al., 2006). Thus, it was proposed that the three WRKY proteins function redundantly as negative regulators of SA-dependent pathways but play a positive role in JA-mediated pathways. Arabidopsis WRKY40 has been shown to bind to the promoter region of *JAZ8* and the *wrky18 wrky40* double mutant constitutively expresses high levels of a number of JAZ family members (Pandey et al., 2010). *Ca*WRKY40a binds to a promoter region of the *JAZ8* ortholog from pepper containing two W-boxes and *CaWRKY40* silenced plants show increased expression of *JAZ8* upon infection with *Xcv* accompanied by a loss of JA-associated leaf phenotypes. Thus, similar to the findings in Arabidopsis the data suggest that *Ca*WRKY40a positively affects JA signaling by directly reducing the expression of at least one of its negative regulators. The phenotype of pepper plants infected with an *XcvΔXops* strain implies that XopS enhances the repressor activity of WRKY40 as these plants show less leaf chloroses compared to wildtype *Xcv* infected plants. In turn, either translocation of XopS during *Xcv* infection or ectopic expression of the effector protein in transgenic Arabidopsis lines lead to an induction of JA-mediated responses. This is in line with the observed effect of XopS on stomatal movement where an induction of JA-signaling would result in an increased stomatal aperture in response to a MAMP stimulus (Melotto et al., 2006). Collectively our data suggest that by targeting WRKY40, XopS positively affects the repressor function of the TF to dampen SA-responses and at the same time induce JA-responses.

Targeting of WRKY TFs by bacterial T3Es is not without precedence. The *Ralstonia solanacearum* effector protein PopP2 is an acetyltransferase that binds and acetylates a range of Arabidopsis WRKY proteins (Le Roux et al., 2015; Sarris et al., 2015). PopP2 acetylation within the WRKY domain of multiple defense-acting WRKY TFs interferes with their binding to W-box containing promoter elements, thereby reducing defense gene activation (Le Roux et al., 2015). Intriguingly, group IIa WRKY TFs *At*WRKY40 and *At*WRKY60 do not interact with PopP2 and are no acetylation substrate for the effector protein. It has been suggested that PopP2 has evolved a degree of substrate discrimination, which avoids acetylation of negative regulators of defense whose inactivation would be disadvantageous for bacterial infection (Le Roux et al., 2015). Our data imply that binding of XopS to WRKY TFs has the opposite selectivity relative to PopP2. XopS interacts with WRKY40 orthologs from different plants but does not bind to *Nb*WRKY8, a group I WRKY TF, which promotes defense gene induction (Ishihama et al., 2011). Similar to the effect of acetylation of negative regulators of defense gene expression by PopP2, stabilization of positive regulators by XopS would counteract the virulence activity of the effector.

Thus, stabilization of negatively acting group IIa WRKY TFs, such as WRKY40, by XopS represents a novel mechanism to reduce defense gene induction and at the same time induce JA-signaling, providing an additional means to counteract SA-mediated immunity.

The mechanism through which XopS stabilizes the WRKY40 protein is currently unknown. The available data suggest that WRKY40 is subject to proteasomal turnover likely as a means to balance its repressor activity during active defense and that XopS interferes with degradation of the TF by the proteasome. Several other WRKY TFs have been described to be degraded by the proteasome to regulate expression of their target genes in response to a range of stimuli, including immunity (Miao and Zentgraf, 2010; Matsushita et al., 2013; Ye et al., 2018; Liu et al., 2021). E3-ubiquitin ligases from different families were shown to be involved in WRKY ubiquitination. Recently, the *Phytophthora sojae* effector protein Avr1d was demonstrated to repress the E3-ligase activity of *Gm*PUB13 to facilitate infection (Lin et al., 2021). However, it seems unlikely that XopS acts as inhibitor of the E3-ligase catalyzing WRKY40 ubiquitination as this would not require a direct interaction with the TF and moreover our data suggest that XopS stabilized WRKY40 is still conjugated to ubiquitin indicative of modification by an E3-activity. Ubiquitin modifies protein substrates mostly in the form of a K-48 linked polyUb chain, which serves as a signal for proteasome degradation (Finley, 2009). However, other types of ubiquitination, including mono-ubiquitination and poly-ubiquitination, in which ubiquitin chains are formed through the linkage of lysine residues other than K48 of the ubiquitin molecule, i.e. K6-, K11-, K27-, K29-, K33- and K63-linked ubiquitination, or through the C- or N-terminus of the ubiquitin moieties (i.e. linear ubiquitination), may also occur (Kulathu and Komander, 2012). Although not very well understood, many non-K48-linked types of ubiquitination appear to serve as non-proteolytic, regulatory modifications often related to chromatin organization, transcription, DNA repair, and protein trafficking (Mukhopadhyay and Riezman, 2007). It is currently unclear how these atypical ubiquitin conjugates are assembled or whether XopS influences the ubiquitin chain topology of WRKY40 either directly or by recruiting auxiliary proteins to the XopS/WRKY40 complex.

In conclusion, the data discussed above lead us to propose a working model of how XopS promotes bacterial virulence (Supplementary Figure S14). Future experiments will have to investigate the biological mechanism of how XopS prevents proteasomal degradation of WRKY40 and reveal possible enzymatic activities of the effector. It will be interesting to see if the mechanism of stabilizing host transcription factors to subvert defense responses is widely used by pathogens.

## Material and Methods

### Plant material and growth conditions

Pepper (*Capsicum annuum* cv. Early Cal Wonder (ECW)) and tobacco plants (*Nicotiana benthamiana*) were grown in soil in a growth chamber with daily watering, and subjected to a 16 h light : 8 h dark cycle (25°C : 20°C) at 240-300 µmol m^−2^ s^−1^ light and 75% relative humidity. Transgenic β-estradiol (Sigma-Aldrich) inducible XopS-GFP Arabidopsis (*Arabidopsis thaliana*) lines were generated by transformation of Col-0 wild type plants with a plasmid containing the entire XopS coding region inserted into binary vector pABindGFP (Bleckmann et al., 2010) using the floral dip method (Clough and Bent, 1998).

For the selection of transgenic plants, seeds of T0 plants were sterilized and sown onto Murashige and Skoog medium (Sigma-Aldrich) supplemented with Gamborg’s vitamin solution (1:1.000) and 20 µg mL^-1^ hygromycin B (Roth). Primary transformants were allowed to self-fertilize and were propagated into the homozygous T3 generation. Arabidopsis plants were grown on soil in an 8 h light : 16 h dark cycle (22°C : 18°C) at 80 µmol m^−2^ s^−1^ light and 70% relative humidity. Transgene expression was induced 6 weeks after germination by spraying with 50 µM β-estradiol and 0.1% Tween-20. Control plants were sprayed with 0.1% EtOH in water and 0.1% Tween-20.

### Plasmid construction

To generate plasmids containing the corresponding gene of interest, the entire open reading frame either with or without the stop codon, was amplified from cDNA. The resulting fragments were inserted into the pENTR-D/TOPO vector according to the manufacturer’s instructions (ThermoFisher) and verified by sequencing. For Y2H experiments, fragments containing a stop codon were recombined into Gateway-compatible versions of the GAL4 DNA BD vector pGBT-9 and the activation domain vector pGAD424 (Clontech) using L/R-clonase (ThermoFisher). To generate translational fusions between the protein of interest and Green fluorescent protein (GFP), coding sequences without stop codon were recombined as described above into the binary vector pK7FWG2 (Karimi et al., 2002). Constructs for bimolecular complementation analysis are based on Gateway-cloning-compatible versions of pRB-C-Venus^N173^ and pRB-C-Venus^C155^. To tag proteins with Hemagglutinin (HA), fragments were inserted into pGWB614 (Nakamura et al., 2010). Constructs for protein expression in *Escherichia coli* (*E. coli*) were either inserted into a Gateway-cloning-compatible version of pMal-C2 (New England Biolabs, NEB) to create a fusion to the Maltose-binding protein (MBP) or into pDEST15 for fusions to Glutathion-S-transferase (GST) (ThermoFisher). To create GUS reporter constructs, either an oligonucleotide containing four W-boxes and the minimal cauliflower mosaic virus (CaMV) 35S promoter, or the 2 kb region upstream of the translational start site of *CaPR4* were inserted in front of the GUS reporter gene in vector pRB-bar-GUS using the NEBuilder recombinational cloning kit (NEB).

Constructs for virus-induced gene silencing in pepper or *N. benthamiana* were generated by recombining a PCR fragment of the gene of interest from pENTR-D/TOPO into pTRV2-Gateway (Liu et al., 2002).

Sequences of oligonucleotides used for cloning are available upon request.

### Yeast two-hybrid analysis

Yeast two-hybrid techniques were performed according to the yeast protocols handbook and the Matchmaker GAL4 Two-hybrid System 3 manual (both Clontech, Heidelberg, Germany) using the yeast reporter strains AH109 and Y187. To identify XopS interacting proteins, a GAL4 AD-domain tagged cDNA library from *Nicotiana tabacum* (Börnke, 2005) was screened as detailed in Üstün et al., (2013a).

### Yeast transactivation assay

WRKY proteins used for this experiment were fused into pGBT9 vector (containing GAL4 DNA-BD) and transformed into yeast strain Y190. Transformed clones were selected on SCAD yeast growth medium (6.7 g/L yeast nitrogen base, 20 g/L glucose, 0.67 g/L amino acid mix, 200 mg/L adenine hemisulfate; pH 5.8) lacking Trp (-T) and growing cells were spotted onto selective medium lacking Trp and His (-TH), supplemented with 10 mM 3-amino-1,2,4-triazole (3-AT). Three days later, *HIS3* reporter gene activity was monitored by the ability of transformed cells to grow on SCAD –TH plates. Additionally, cells growing on selective medium were further tested for their ability to induce the *lacZ* reporter gene using a filter lift assay. The empty pGBT9 vector was used as negative control and the pGAL4 vector containing full length GAL4 was used as positive control. Yeast cells were grown at 30°C.

### Bacterial strains and plant infection assays

Strains used in this study were as follows: *Xanthomonas campestris* pv. *vesicatoria* (*Xcv* wild type), *XcvΔhrpF* (type-3 secretion-deficient mutant), *XcvΔxopS,* complementation strain *XcvΔxopS*/*XopS-HA* and *Pseudomonas syringae* pv. *tomato* strain DC3000 *PstΔhrcC* (type-3 secretion-deficient mutant). The *XcvΔxopS* deletion strain and its *XcvΔxopS*/*XopS-HA* complementation strain were constructed as described in Üstün et al. (2013b) using the primers listed in Supplementary Table S1. Bacteria were grown overnight in KingsB medium (20g/L glycerol, 40g/L peptone, 10% K_2_HPO_4_, 10% MgSO_4_) containing appropriate antibiotics at 28°C with shaking. For *in planta* growth assays using syringe-inoculation, leaves from four-to five-week-old pepper plants were infiltrated with the different *Xcv* strains at OD_600_ = 0.0001 (1×10^5^ CFU mL^-1^) using a 1 mL sterile needleless syringe. Leaves from six-week-old Arabidopsis plants were inoculated with *PstΔhrcC* at OD_600_ = 0.00002 (1×10^4^ CFU mL^-1^). Bacterial suspensions were prepared in 10 mM MgCl_2_. For dip-inoculation assays of pepper plants with *Xcv* strains, bacterial suspensions were prepared at OD_600_ = 0.2 (1×10^8^ CFU mL^-1^) in 10 mM MgCl_2_ supplemented with 0.02% (v/v) silwet l-77 (Helena Chemical Company). One leaf per plant (four-to five-week-old) was completely submerged into the indicated bacterial suspension for 1 min. Leaves were then covered with a moisturized plastic bag for 24 h to ensure 100% humidity. Plants were kept in the growth chamber under a 16 h light: 8 h dark cycle (25°C : 20°C) at 240-300 µmol m^−2^ s^−1^ light and 75% relative humidity. For determination of bacterial titer of each analysed strain, one or two leaf discs (1 × 0.5 cm^2^ for pepper, 2 × 0.22 cm^2^ for Arabidopsis) per biological replicate were collected, pooled and ground in 10 mM sterile MgCl_2_. Two technical replicates per biological replicate were prepared. Serial dilutions were spotted onto KingsB plates supplemented with appropriate antibiotics. For infection assays used for analysis of marker gene expression, leaves from four-to five-week-old pepper plants were infiltrated with the different *Xcv* strains at OD_600_ = 0.2 (1×10^8^ CFU mL^-1^) in 10 mM MgCl_2_. 10 mM MgCl_2_ was used as mock control. At least three biological replicates were used per experiment unless otherwise stated.

### Transient expression of proteins in N. benthamiana

For infiltration of *N. benthamiana* leaves, *Agrobacterium tumefaciens* (*Agrobacterium*) strain C58C1 was infiltrated into the abaxial air space of four-week-old plants, using a 1 mL sterile needleless syringe. *Agrobacteria* were cultivated overnight in YEB medium (5g/L Difco bovine extract, 1g/L yeast extract, 1g/L bacto peptone, 5g/L saccharose, 2mM MgSO_4_) containing appropriate antibiotics at 28°C with shaking. The cultures were harvested by centrifugation, and the pellet was resuspended in infiltration buffer (10 mM MgCl_2_, 10mM MES pH 5.7, 200 µM acetosyringone; Sigma-Aldrich) and incubated for two hours in the dark with shaking prior to infiltration. Cultures containing different transgenes were infiltrated at OD_600_ = 0.5.

*Agrobacteria* carrying the silencing inhibitor P19 were co-infiltrated at OD_600_ = 0.3. Infiltrated plants were kept in the growth chamber under 16 h light: 8 h dark cycle (25°C : 20°C) at 240-300 µmol m^−2^ s^−1^ light and 75% relative humidity.

### Virus-induced gene silencing (VIGS)

VIGS was performed as described previously (Üstün et al., 2013). Briefly, *Agrobacterium* strains carrying the pTRV1 vector (Liu et al., 2002) and pTRV2-*GFP* or pTRV2-*CaWRKY40a* (OD_600_ = 1.0) were mixed in a 1:1 ratio, respectively, and the mixture was infiltrated into cotyledons of 10-day-old pepper plants using a 1 mL sterile syringe without a needle. For each experiment, 10 plants were infiltrated with pTRV2-*GFP* (control) and 50 plants were infiltrated with pTRV2-*CaWRKY40a*. For VIGS in *N. benthamiana*, *Agrobacterium* strains carrying the pTRV1 vector and pTRV2 empty vector (pTRV2) or pTRV2-*NbWRKY40* (OD_600_ = 1.0) were mixed in a 1:1 ratio, respectively, and the mixture was infiltrated into two lower leaves of two-week-old *N. benthamiana* plants (four-leaf-stage) using a 1 mL sterile needleless syringe. For each experiment, 10 plants were infiltrated with pTRV2 (control) and 30 plants were infiltrated with pTRV2-*NbWRKY40*. The *Agrobacterium*-inoculated plants were kept in dark for 56 h and then grown under a 16 h light: 8 h dark cycle (25°C : 20°C) at 240-300 µmol m^−2^ s^−1^ light and 75% relative humidity. Silenced plants were analysed two-to three-weeks post inoculation. The mRNA levels of silenced plants were checked via quantitative real-time PCR upon treatment of one excised leaf per plant with 5 mM salicylic acid (SA) for 4 h to trigger *WRKY40* gene expression. *WRKY40* mRNA levels in pTRV2-*CaWRKY40a* pepper or pTRV2-*NbWRKY40 N. benthamiana* plants were compared to mRNA levels in pTRV2-GFP or pTRV2 plants, respectively.

### RNA extraction and quantitative real-time PCR

Total RNA was isolated from leaf material and then treated with RNase-free DNase to degrade any remaining DNA. RNA concentrations were measured using a microplate reader (Tecan Infinite 200 PRO). First-strand cDNA synthesis was performed from 1 µg of total RNA using Revert-Aid reverse transcriptase (ThermoFisher). For quantitative real-time PCR (qRT-PCR), the cDNAs were amplified using SensiFAST SYBR Lo-ROX Mix (Bioline) in the AriaMx Realtime PCR System (Agilent Technologies) as previously described (Arsova et al., 2010). At least three biological and two technical replicates were used for each analysis.

The transcript level was standardized based on cDNA amplification of the reference genes stated in the respective figure legends. Primer sequences are provided in Supplementary Table S1.

### Stomatal assays

Leaf discs were cut from leaves of four-to five-week-old pepper and *N. benthamiana* plants or from six-week-old Arabidopsis plants and floated on water for 2 h to allow maximum stomatal opening. After 2 h, *N. benthamiana* or Arabidopsis leaf discs were floated on water supplemented with 25 µM flg22 (Hycultec) or 50 µM abscisic acid (ABA; Sigma-Aldrich) for 2 h as described in Lozano-Durán et al. (2014). Pepper leaf discs were floated on water supplemented with a bacterial suspension of different *Xcv* strains at OD_600_ = 0.2 (1×10^8^ CFU mL^-1^) for 2 h.

Leaf discs were kept under light (240-300 µmol m^−2^ s^−1^) over the whole duration of the experiment and were removed from the growth chamber only right before the microscopic analysis. At the indicated time points, leaf discs were removed to a microscope slide containing sterilized water and the lower epidermis of the leaf discs was observed under a digital microscope VHX-7000 (Keyence, version 1.4.17.3). For each experiment, stomatal apertures (width/length) of approximately 100 stomata per treatment were measured using the onboard software of the microscope. Completely closed stomata were reported as a value of 0 μm. At least three biological replicates were used per experiment unless otherwise stated.

### Chlorophyll measurement

For determination of total chlorophyll (Chl *a* + *b*) content, four leaf discs were cut from each biological replicate and incubated in 1 mL of 80% acetone overnight at 4°C with shaking. Measurements were made by pipetting 200 µL sample into a 96-well flat bottom polystyrene plate (Sarstedt) and absorption at wavelength 663 nm (Chl *a*) and wavelength 645 nm (Chl *b*) was read in a Tecan Infinite 200 PRO microplate reader. 80% acetone was used as blank. Total chlorophyll was calculated using the formula described in Lichtenthaler (1987). Chlorophyll content in control plants was set to 100%. Two technical replicates per biological replicate were measured. At least six biological replicates were used per experiment unless otherwise stated.

### Plant hormone analysis

Salicylic acid (SA) was extracted from frozen leaf material and analyzed by ultra-high-performance liquid chromatography system (Agilent Technologies) coupled to an Agilent 6530 quadrupole time of flight-mass spectrometer (Agilent Technologies) using the internal standard d4-SA, as described previously (Nietzsche et al., 2018). Jasmonic acid (JA) was extracted from the plant matrix and analyzed by high-performance liquid chromatography (HPLC) coupled to a QTrap® mass spectrometer. In brief, 5 mg of freeze-dried and homogenized sample material was extracted with a methanol/water (60:40, v/v) solution. Deuterated jasmonic acid (d5-JA) was added as internal standard. Analysis was performed using an Agilent Technologies 1260 Infinity HPLC coupled to Q-Trap®6500 ESI-MS/MS system (Sciex, USA). The analytes were separated on an Eclipse Plus C18 column (Agilent Technologies, Waldbronn, Germany) using water (+0.1 % acetic acid) and acetonitrile (+0.1 % ultrapure water) as a mobile phase in a gradient mode.

Ionization was performed with electron spray ionization (ion source temperature, 500°C; ion spray voltage, −4,500 V; curtain gas, 50 psi; nebulizer gas, 50 psi; auxiliary gas, 65 psi) in the negative ionization mode. Multiple reaction monitoring (MRM) mode was used for the measurement, and the transitions of JA (209 → 59; collision energy, −16 V, entrance potential −5 V; cell exit potential −15 V; declustering potential −50 V) and d5-JA (214 → 62; collision energy −16 V, entrance potential −5 V; cell exit potential −15 V; declustering potential −50 V) were used for quantification. The concentrations were calculated based on external calibration curves of JA and the recovery of the internal standard (d5-JA).

### Immunoblotting

Transiently expressed proteins were extracted from *N. benthamiana* leaf material using plant protein extraction buffer (50 mM Tris-HCl pH 7.5, 150 mM NaCl, 5 mM EDTA, 1 mM NaF, 10 mM DTT, 0.1% Triton X-100 and 1% [v/v] protease inhibitor cocktail; Sigma-Aldrich). Protein extracts were boiled at 95°C for 10 min in 1x sodium-dodecyl sulfate (SDS) loading buffer (4x; 200 mM Tris-HCl pH 6.8, 0.4 M DTT, 40% glycerol, 6 mM bromphenol blue and 8% SDS) and then subjected to SDS-polyacrylamide gel electrophoresis (SDS-PAGE). Separated proteins were transferred onto nitrocellulose membrane (Amersham), blocked with 5% skimmed milk in TBS-T and incubated with respective antibodies in 1% skimmed milk in TBS-T. Antibodies used in this study were as following: horseradish peroxidase-conjugated anti-GFP (1:1.000, SantaCruz), anti-HA (1:500, Sigma-Aldrich) and anti-Ubiquitin (1:500, SantaCruz). Proteins were detected using the Clarity Western ECL substrate (BioRad) and chemiluminescence was detected using a ChemiDoc imager (Bio Rad).

### Bimolecular fluorescence complementation (BiFC)

Constructs were transformed into *Agrobacterium* strain C58C1 and transiently expressed by *Agrobacterium*-infiltration in *N. benthamiana*. The BiFC-induced YFP fluorescence was detected by CLSM (LSM510; Zeiss) 48 hours post inoculation (hpi). The specimens were examined using the LD LCI Plan-Apochromat 253/0.8 water-immersion objective for detailed images with excitation using the argon laser (458- or 488-nm line for BiFC and chlorophyll autofluorescence). The emitted light passed the primary beamsplitting mirrors at 458/514 nm and was separated by a secondary beam splitter at 515 nm. Fluorescence was detected with filter sets as follows: on channel 3, 530–560 band pass; and on channel 1, for red autofluorescence of chlorophyll.

### In planta GFP pull-down

Approximately 2 g of leaf material was ground to a fine powder in liquid nitrogen and homogenized in 4 mL of extraction buffer (25 mM Tris-HCl, pH 7.5, 150 mM NaCl, 10%glycerol, 1 mM DTT, 1 mM EDTA, 0.5% Triton X-100 and 1% [v/v] protease inhibitor cocktail; Sigma-Aldrich). Insoluble debris was pelleted by centrifugation for 20 min with 4000 rpm at 4°C. Immunoprecipitation was performed by adding 30 µL of GFP-Trap^®^ coupled to magnetic beads (ChromoTek) and samples were incubated for 2 h at 4°C with continual rotation. Beads were subsequently washed five times with Tris-buffered saline containing 0.5% Triton X-100, and immunoprecipitates were eluted with 30 μL of 2x SDS loading buffer at 70°C for 10 min.

### In vitro pull-down

Recombinant MBP-*Nb*WRKY40 or MBP alone from *E. coli* lysates were immobilized on amylose resins (NEB), incubated for 1 h at 4 °C with purified GST-XopS, eluted, and analyzed by immunoblotting using either anti-GST antibody (1:1.000, horseradish peroxidase-conjugated; Sigma-Aldrich) or anti-MBP antibody (1:10.000, used with 1:10.000 secondary horseradish peroxidase-conjugated anti-mouse antibody; NEB).

### GUS activity measurement

For *in planta* GUS reporter assays, four-week-old *N. benthamiana* plants were inoculated with *Agrobacteria* carrying indicated effector constructs, alongside with *Agrobacteria* carrying indicated reporter constructs at OD_600_ = 0.5. *Agrobacteria* carrying the silencing inhibitor P19 were co-infiltrated at OD_600_ = 0.3. After 48 h, four leaf discs (0.5 cm^2^) per sample were ground in 120 µL GUS extraction buffer (50 mM sodium phosphate buffer pH 7.0, 10 mM DTT, 1 mM Na_2_EDTA, 0.1% SDS and 0.1% Triton X-100), incubated for 20 min on ice and then centrifuged at 13.000 rpm for 10 min at 4°C. Protein concentrations were determined using Bradford reagent (BioRad) and 200 µg of protein extract were pre-incubated in 450 µL GUS assay buffer for 10 min at 22°C, to guarantee equal protein amounts incubated with the substrate. The reaction was started by adding 25 µl of 20 mM 4-Methylumbelliferyl-β-D-glucuronide hydrate (MUG; Sigma-Aldrich) substrate to the samples and a 20 µl aliquot was immediately added to 180 µl stop buffer (0.2 M Na_2_CO_3_) which was used as timepoint 0 control. The rest of the mixture was incubated at 37°C for 1 h and the reaction was stopped by adding again 20 µl aliquots to 180 µl stop buffer. GUS activity was determined by measuring the fluorescence (excitation wavelength 365 nm and emission wavelength 460 nm) of produced 4-Methylumbelliferon (4-MU) in a 96well polystyrene microwell plate (Nunc F96 black flat bottom, ThermoFisher) by the Tecan Infinite 200 PRO microplate reader. A 4-MU standard curve was used to calculate the produced amount of 4-MU per unit of time and GUS activity is given in pmol 4-MU min^-1^ mg protein^-1^. Protein expression of effectors was verified by immunoblotting.

### Fluorescent electro mobility shift assay

DNA binding reactions for the fluorescent electrophoretic mobility shift assay were carried out in 25 mM HEPES-KOH pH 7.6, 40 mM KCl, 1 mM DTT and 10% Glycerol. 50 ng Cy5-labeled probe DNA were incubated with indicated recombinant proteins in a total volume of 20 μL for 30 min at room temperature. MBP-*Ca*WRKY40a and MBP recombinant proteins were produced in *E. coli* and purified using amylose resin (NEB). Target probes for MBP-*Ca*WRKY40a upstream regions consisted of 150 bp fragments containing two predicted W-boxes that were amplified by PCR using Cy5-labelled primers and gel purified. As competitor DNA, the same fragments amplified using unmodified primers were used and added in 50-fold (50x) excess over the labelled probe. Primer sequences can be found in Supplementary Table S1. 1x Orange-G (0.25% Orange-G and 30% glycerol) was used as loading dye. The DNA–protein complexes were resolved on 5 % non-denaturing polyacrylamide gels (Bio Rad) and visualized using an Octoplus Q-Plex Imaging System (Intas).

### Drug treatment

For analysing protein stability in *N. benthamiana* plants transiently expressing binary *Nb*WRKY40 constructs, leaves were infiltrated 42 hpi with 200 µM MG132 (Sigma-Aldrich). Six hours later, leaf material was harvested and processed.

### Proteasome activity measurement

The experiment was performed as previously described (Üstün and Börnke, 2017). Proteasome activity is expressed in percentage relative to the control treatment (set to 100%).

### Multiple sequence alignment

An alignment of different WRKY proteins was obtained using Clustal W with default parameters (www.ebi.ac.uk). Shadings were made with BOXSHADE 3.21 (https://embnet.vital-it.ch/software/BOX_form.html).

### Statistical analysis and data presentation

Depending on the experiment, statistical significances were based on either Student’s *t*-test or one-way ANOVA followed by Tukey’s multiple comparisons test. The model generated to elucidate XopS mode of action was created with BioRender.com. Each experiment was repeated at least twice unless otherwise stated. The number of biological replicates (individual plants, n) is stated in the figure legends.

### Accession numbers

Sequence data for genes relevant to this article can be found in the Arabidopsis Genome Initiative or GenBank/EMBL databases under the following accession numbers: *AtWRKY40*, AT1G80840; *CaWRKY40*; XP_016578457; *CaWRKY40a*, XP_016562883, *NtWRKY40*, XP_016457007; *NbWRKY40*; Niben101Scf06091g04005.1; *NbWRKY8*, AB445392, *CaCDPK15*, XM_016716796.1; *CaPR1*, AY560589.1; *CaPR4*, JX030397.1, *CaJAZ8*, XM_016682106.1 and *XopS*, CAJ21955.

## Acknowledgements

We thank Kerstin Bieler for her skillful technical help. This study was supported by funds from the Deutsche Forschungsgemeinschaft (DFG; German Research Foundation) to F.B. (BO1916/5-2 and CRC973 project C6). S.Ü. was supported by an Emmy Noether Fellowship GZ: UE188/2-1 awarded by the DFG.

## Author contributions

M.R., S.Ü., and F.B. conceived and designed the experiments. M.R., S.Ü., T.G. D.S. M.F. and F.B. performed the experiments. M.R., S.Ü., S.B., and F.B. analyzed the data. S.S. provided unpublished materials. M.R. and F.B. wrote the manuscript with input from all authors.

## Supplemental Data

**Supplementary Figure S1.**
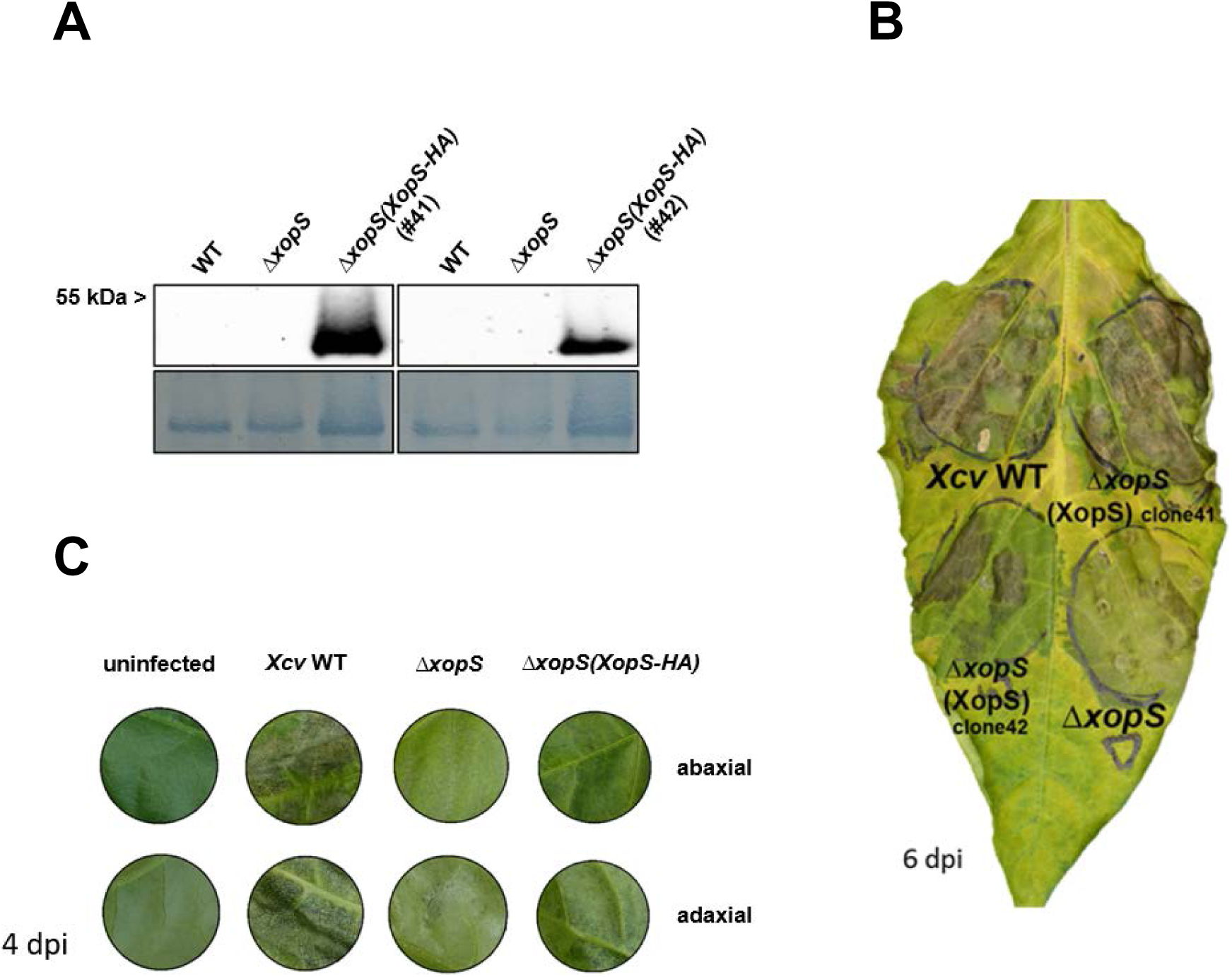
XopS contributes to *Xcv* symptom development on susceptible pepper plants. **(A)** Verification of XopS-HA protein expression in an *XcvΔxopS*(*XopS-HA*) complementation strain. Leaves of susceptible pepper plants (ECW) were syringe-inoculated with *Xcv* wild type*, XcvΔxopS,* or two independent clones of the complementation strain *XcvΔxopS*(*XopS-HA*) at OD_600_ = 0.2. Total protein extracts from pepper leaves were prepared at 3 dpi and equal volumes representing approximately equal protein amounts of each extract were immunoblotted. Proteins were detected using an anti-HA antibody and Amido black staining of RubisCo served as loading control. **(B)** Phenotype of a representative pepper leaf 6 dpi with *Xcv, XcvΔxopS,* or *XcvΔxopS* at OD_600_ = 0.2. Two different clones (clone 41 and clone 42) of the complemented *Xcv xopS* deletion were tested. Clone 41 was used for all further experiments. **(C)** Disease symptom development in pepper leaves infected with *Xcv* wild type*, XcvΔxopS,* or *XcvΔxopS*(*XopS-HA*). Indicated strains were syringe-inoculated with an OD_600_ of 0.2 into susceptible leaves of pepper ECW plants and the disease phenotype of one representative plant was photographed at 4 dpi.

**Supplementary Figure S2.**
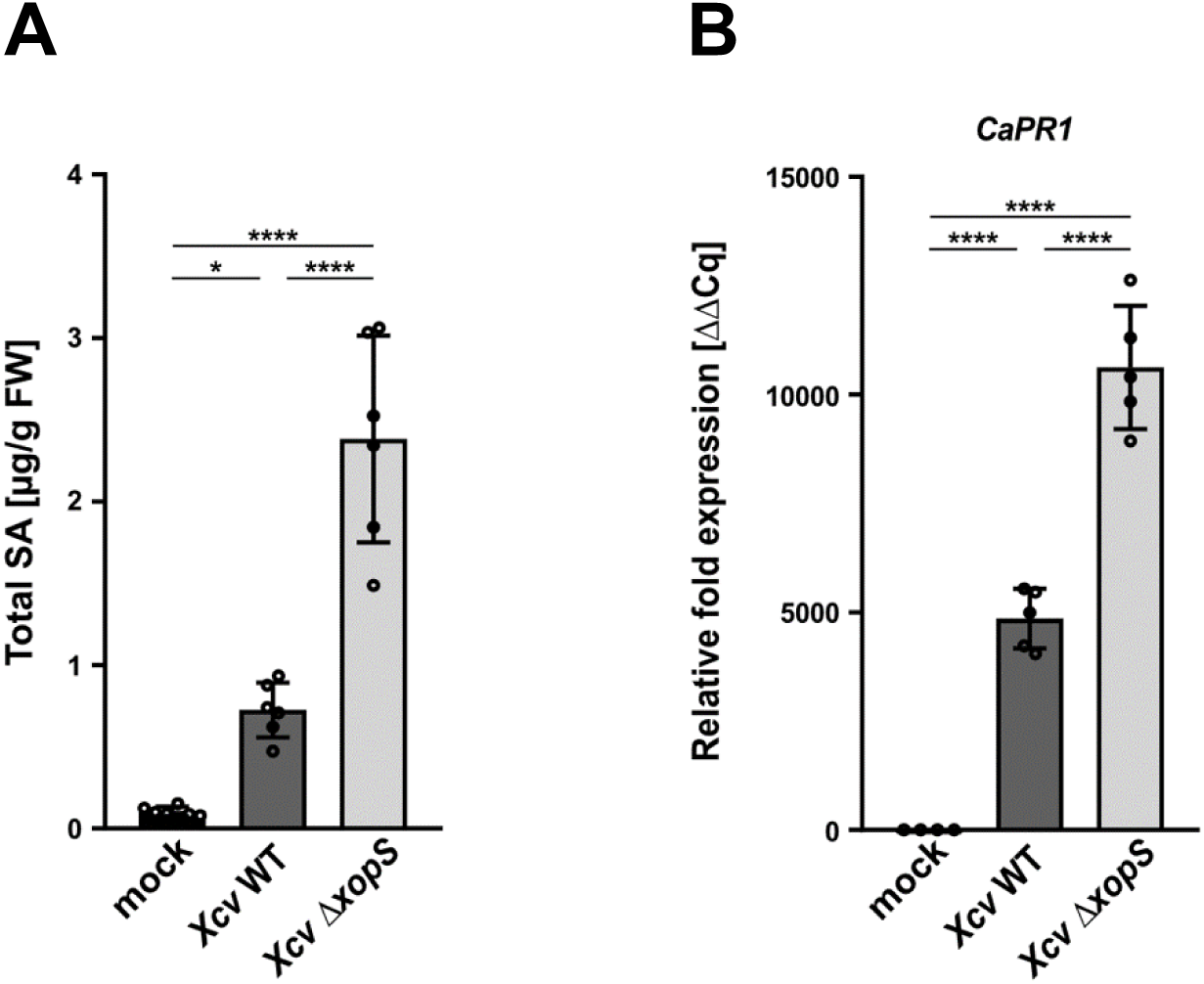
XopS is required for full suppression of SA-dependent defense. **(A)** Salicylic acid (SA) content changes upon infection with different *Xcv* strains in pepper ECW plants. Susceptible pepper leaves were syringe-inoculated with MgCl_2_ (mock) or a bacterial suspension of *Xcv* and *XcvΔxopS* at OD_600_ = 0.2, respectively. Total SA (free SA+SAG) levels in infected tissue were measured 2 dpi. Bars represent the mean of n = 6 biological replicates ± SD and asterisks (*, P < 0.05; ****, P < 0.0001) mark significant differences according to one-way ANOVA. FW, Fresh weight. The experiment was carried out twice with similar results. **(B)** Gene expression analysis of *CaPR1* upon *Xcv* and *XcvΔxopS* infection. Leaves of susceptible pepper plants were syringe-inoculated with indicated strains at OD_600_ = 0.2. Samples were taken 10 hpi, the mRNA level of *CaPR1* was measured by qRT-PCR and compared to mock treated leaves. *Tubulin* was used as a reference gene. Each bar represents the mean of at least n = 4 biological replicates ± SD (n = 4 for mock, n = 5 for *Xcv* and *XcvΔxopS*). Asterisks (****, P < 0.0001) mark significant differences according to one-way ANOVA. *Tubulin* was used as a reference gene. The experiment was carried out at least three times with similar results.

**Supplementary Figure S3.**
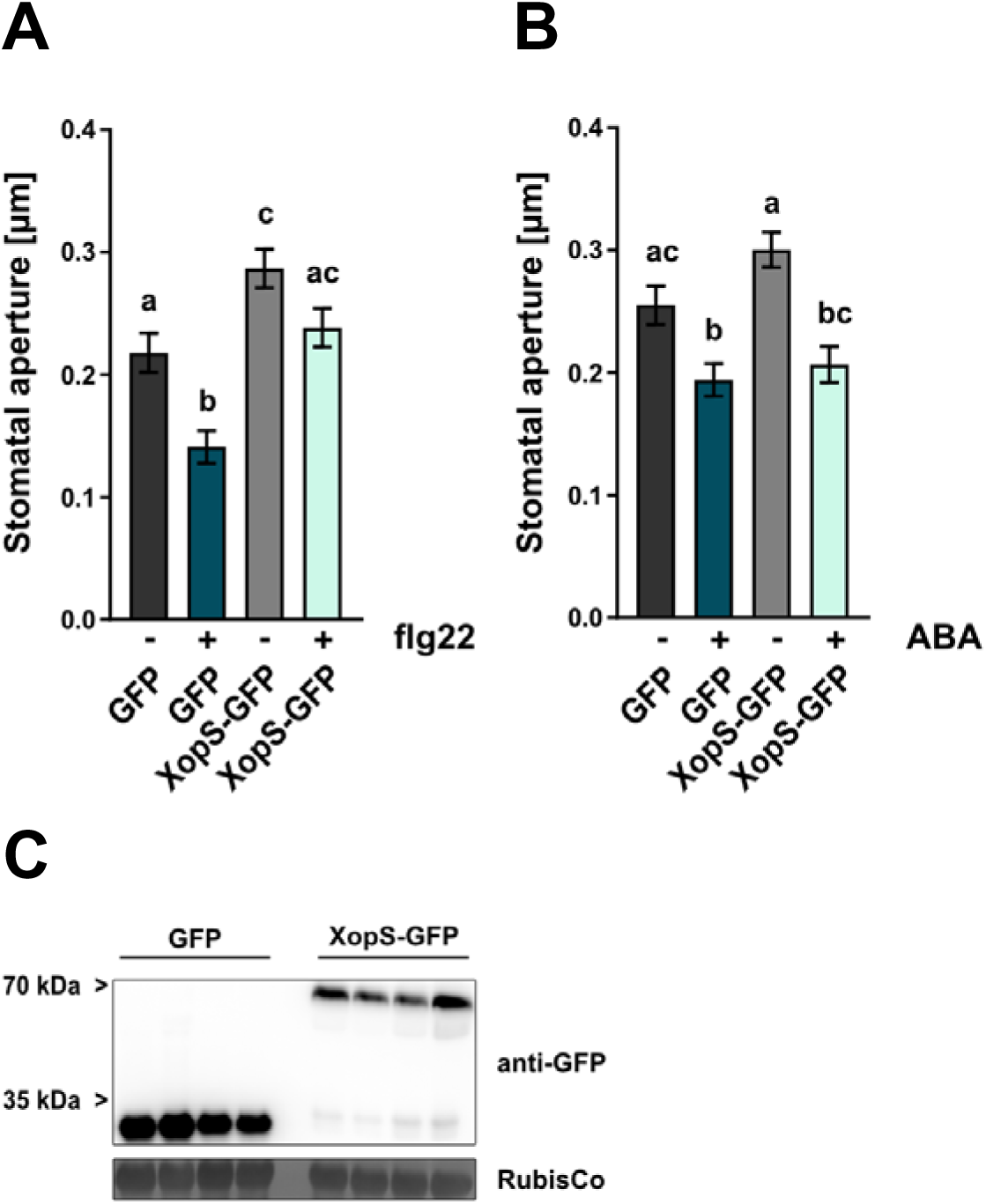
Transient expression of XopS in *N. benthamiana* inhibits stomatal closure in response to flg22. **(A)** Stomatal aperture measurement in *N. benthamiana* plants transiently expressing either free GFP (GFP) or XopS-GFP. Leaf discs were floated on water (control) or on water supplemented with 25 µM flg22 for two hours prior to the measurement of stomatal aperture under a microscope. Approximately 100 apertures from n = 4 independent plants were measured per individual treatment and are represented as width/length ratio. Bars represent the mean ± SE. Letters over bars represent statistical significance determined by one-way ANOVA (P < 0.05). The experiment was carried out twice with similar results. **(B)** Stomatal aperture measurement in *N. benthamiana* plants transiently expressing either GFP or XopS-GFP. Leaf discs were floated on water (control) or on water supplemented with 25 µM flg22 for two hours prior to the measurement of stomatal aperture under a microscope. Approximately 100 apertures from n = 4 independent plants were measured per individual treatment and are represented as width/length ratio. Bars represent the mean ± SE. Letters over bars represent statistical significance determined by one-way ANOVA (P < 0.05). **(C)** Western blot analysis to confirm expression of GFP and XopS-GFP in *Agrobacterium* infiltrated *N. benthamiana* leaves. Samples were taken 24 hpi and proteins were detected using an anti-GFP antibody. Amido black staining of RubisCo served as loading control. The four biological replicates used for the analysis of stomatal aperture are shown.

**Supplementary Figure S4.**
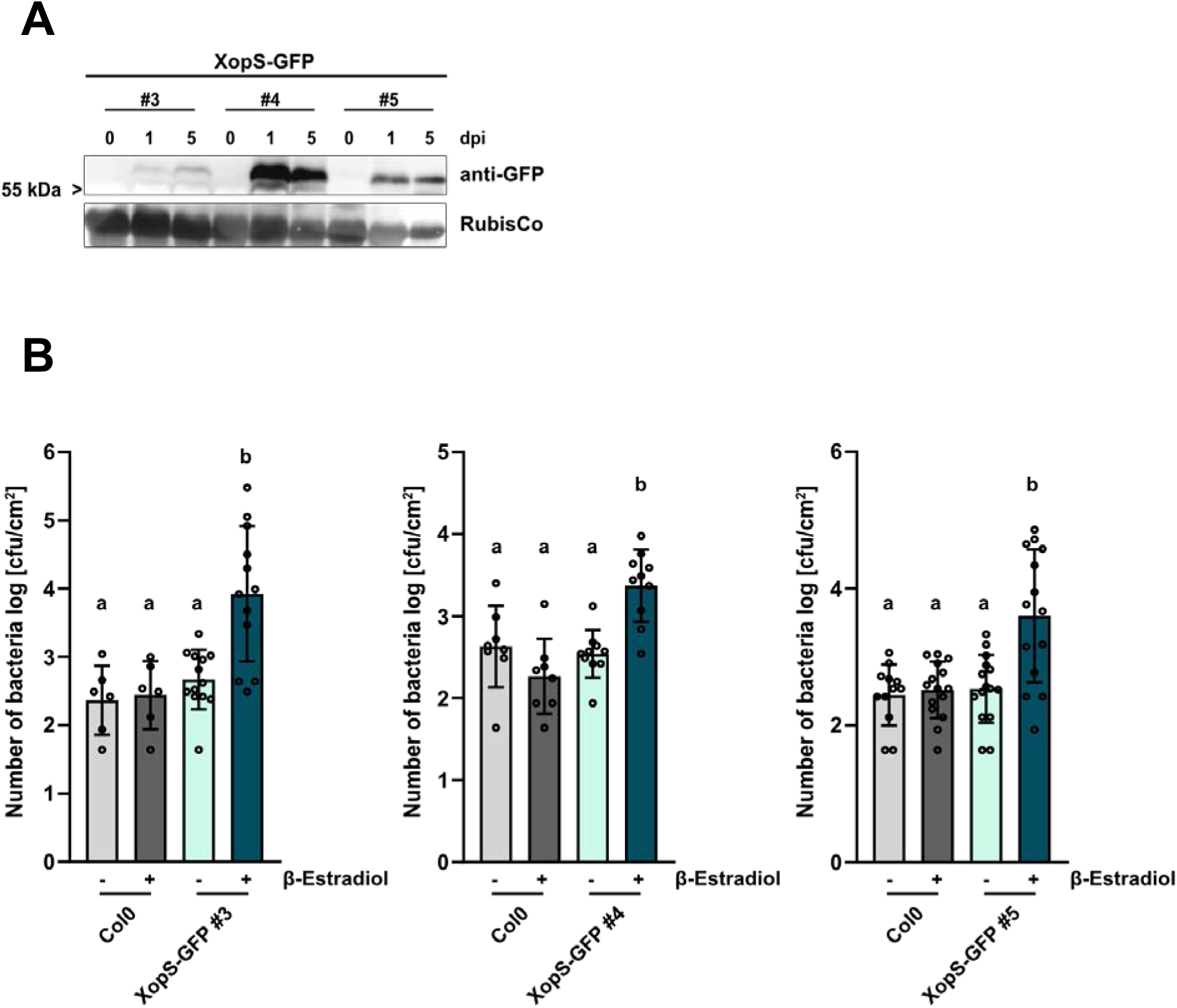
Inducible expression of XopS-GFP in transgenic Arabidopsis lines interferes with PTI. **(A)** Western blot analysis to confirm XopS-GFP protein expression in β-estradiol inducible transgenic *Arabidopsis* lines. 4 weeks old plants grown on soil under short day conditions (8 h light/16 h dark) were sprayed with 50 µM β-estradiol and leaf samples were taken at indicated time points (dpi = days post induction). Expression of the XopS-GFP fusion protein was detected by immunoblotting with anti-GFP antibody. Amido black staining of RubisCo served as loading control. **(B)** Expression of XopS-GFP contributes to the growth of *Pst* DC3000 Δ*hrcC* type-III defective bacteria. Wild-type Col-0 (Col0) and XopS-GFP transgenic plants of indicated independent lines were induced with 50 µM estradiol, syringe-inoculated after 24 h with *Pst* DC3000 Δ*hrcC* bacteria at OD_600_ = 0.00002. Colony forming units were determined 5 dpi. Bars represent the mean of at least n = 3 biological replicates (and 2 technical replicates per biological replicate) ± SD. Letters over bars represent statistical significance determined by one-way ANOVA (P < 0.05). The experiment was carried out three times with similar results.

**Supplementary Figure S5.**
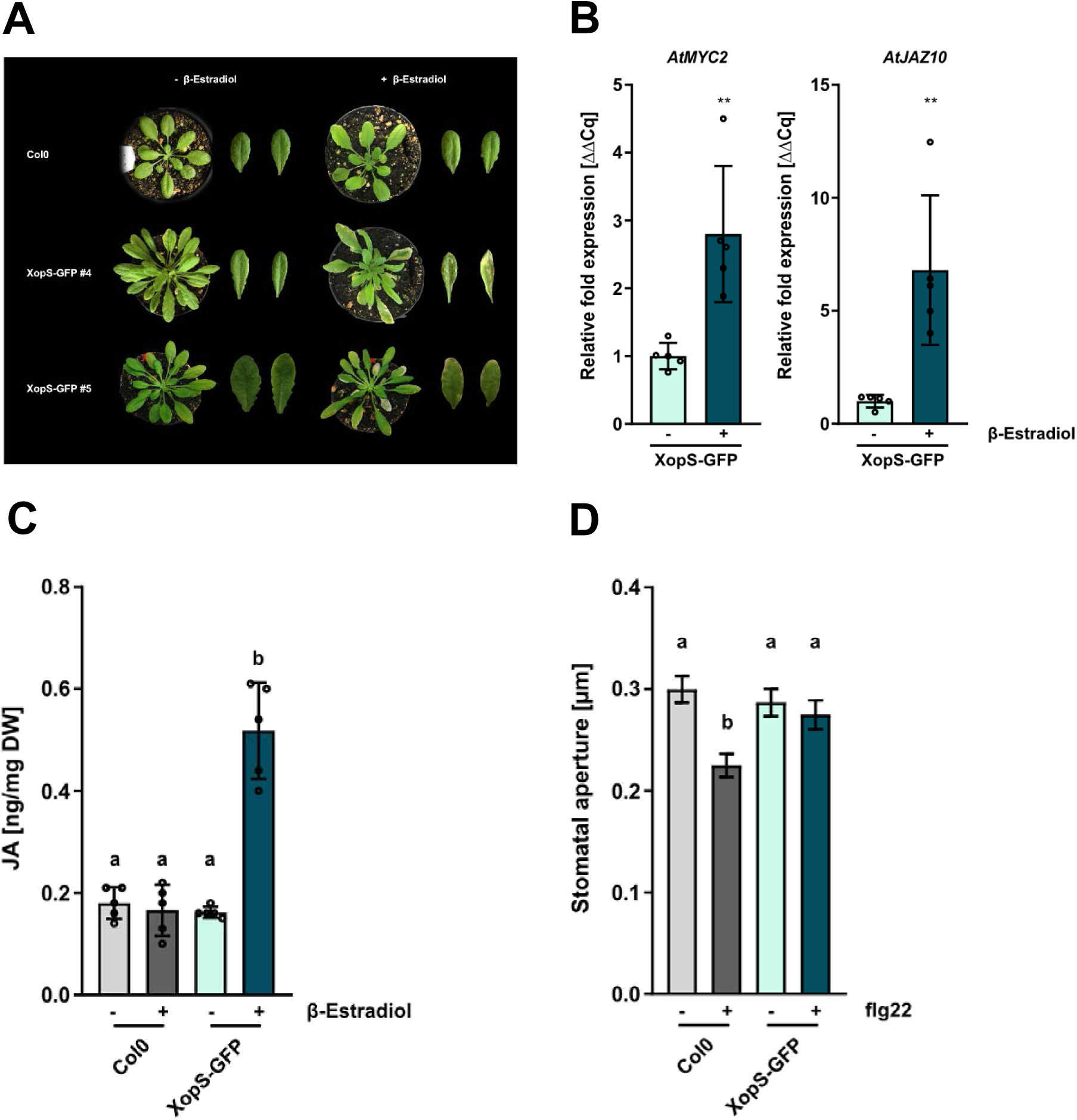
Inducible expression of XopS-GFP triggers JA signaling in Arabidopsis. **(A)** Chlorotic phenotypes of indicated independent transgenic Arabidopsis lines expressing XopS-GFP under the control of the β-estradiol inducible promoter compared to the untransformed wild type Col-0 (Col0) control. Pictures were taken five days after spraying 50 µM β-estradiol. **(B)** Gene expression analysis of JA marker genes in one representative XopS-GFP Arabidopsis line. Samples were taken 4 hours after β-Estradiol induction. Total RNA was extracted and the mRNA levels of indicated marker genes was measured by qRT-PCR and compared to mock treated leaves. *Ubiquitin conjugating enzyme 9* (*UBC9*) was used as a reference gene. Bars represents the mean of n = 5 biological replicates ± SD and asterisks (**, P < 0.01) mark significant differences according to Student’s *t*-test. The experiment was carried out twice with similar results. Jasmonic acid (JA) levels in one representative XopS-GFP transgenic Arabidopsis line 24 hours after induction of protein expression by spraying 50 µM β-estradiol, compared to wild type Col-0 (Col0) plants. - indicates plants that were sprayed with mock control. DW, Dry weight. Bars represent the mean of n = 5 pools of 4 independent plants each ± SD. Letters over bars represent statistical significance determined by one-way ANOVA (P < 0.05). **(D)** Stomatal aperture measurement in wild type Col-0 and transgenic Arabidopsis XopS-GFP lines, with and without induction of protein expression with 50 µM β- estradiol (- and +, respectively). Leaf discs were floated on water (control) or on water supplemented with 25 µM flg22 for two hours prior to the measurement of stomatal aperture under a microscope. Approximately 100 apertures from n = 3 independent plants were measured per individual treatment and are represented as width/length ratio. Bars represent the mean ± SE. Letters over bars represent statistical significance determined by one-way ANOVA (P < 0.05). The experiment was carried out twice with similar results.

**Supplementary Figure S6.**
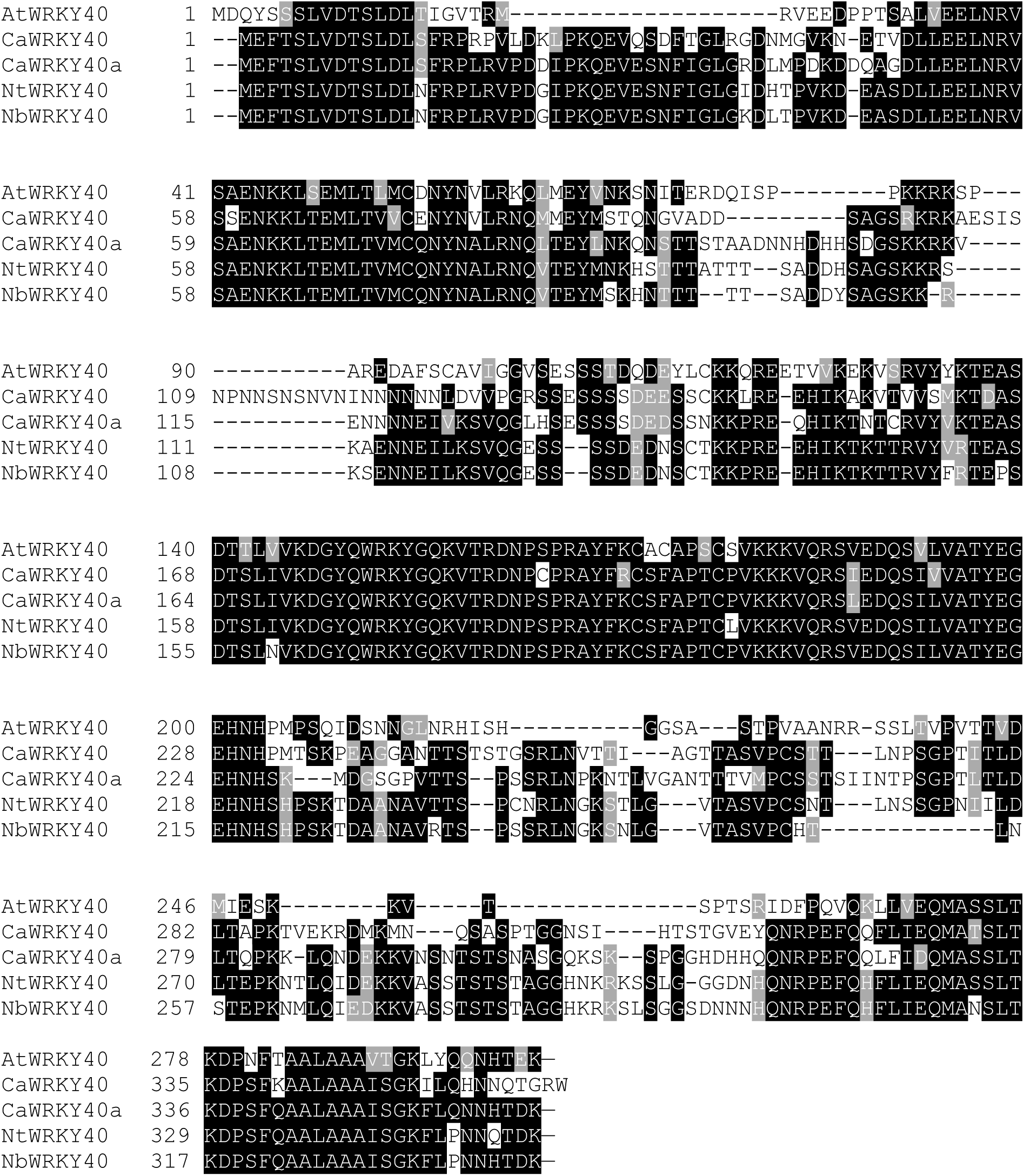
Multiple sequence alignment of WRKY40 proteins. *Arabidopsis thaliana* (*At*WRKY40; AT1G80840), *Capsicum annuum* (*Ca*WRKY40; XP_016578457 and *Ca*WRKY40a; XP_016562883), *Nicotiana tabacum* (*Nt*WRKY40; XP_016457007) and *Nicotiana benthamina* (*Nb*WRKY40; Niben101Scf06091g04005.1) using Clustal W with default parameters (www.ebi.ac.uk). Black and gray shadings, done with BOXSHADE 3.21 (https://embnet.vital-it.ch/software/BOX_form.html), indicate conserved amino acid residues.

**Supplementary Figure S7.**
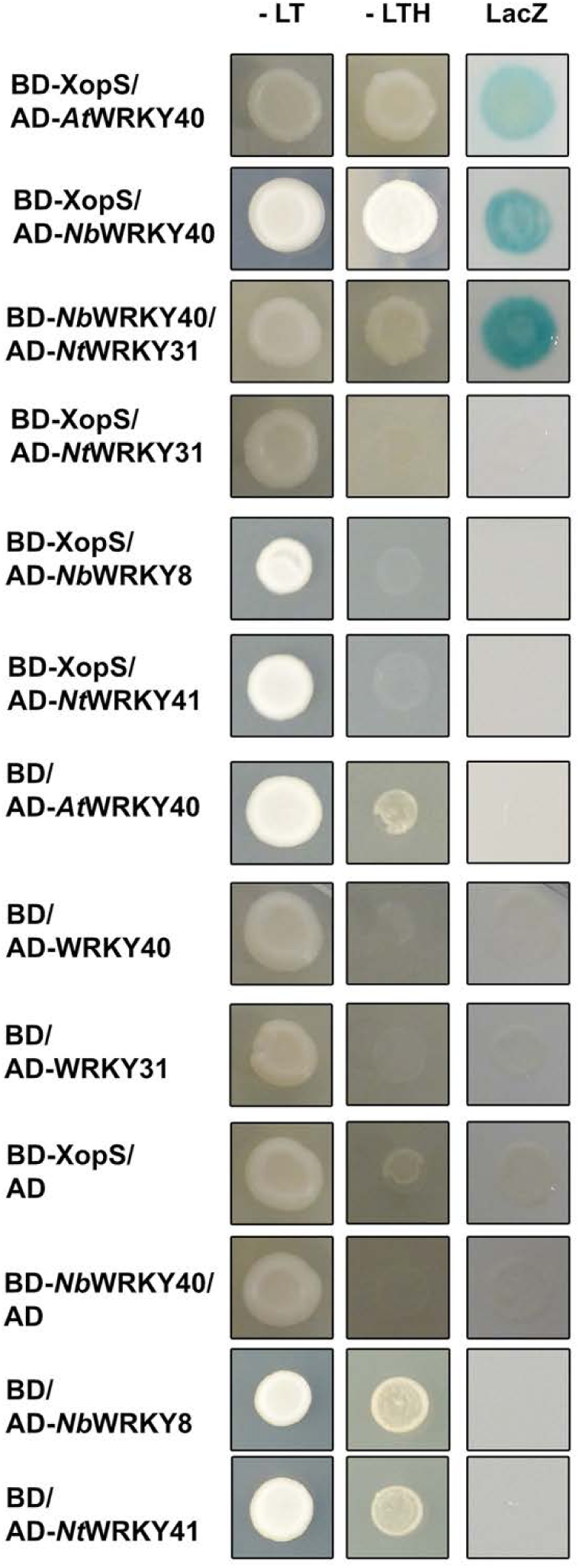
XopS does not interact with WRKY proteins other than WRKY40 in yeast. Proteins were either fused to the GAL4 DNA binding domain (BD) or to the GAL4 activation domain (AD). Indicated combinations were transformed into yeast strain Y190. Cells were grown on selective media before a LacZ filter assay was performed. – LT, yeast growth on medium without Leu and Trp. – LTH, yeast growth on medium lacking Leu, Trp and His indicating expression of the *HIS3* reporter gene. LacZ, activity of the *lacZ* reporter gene.

**Supplementary Figure S8.**
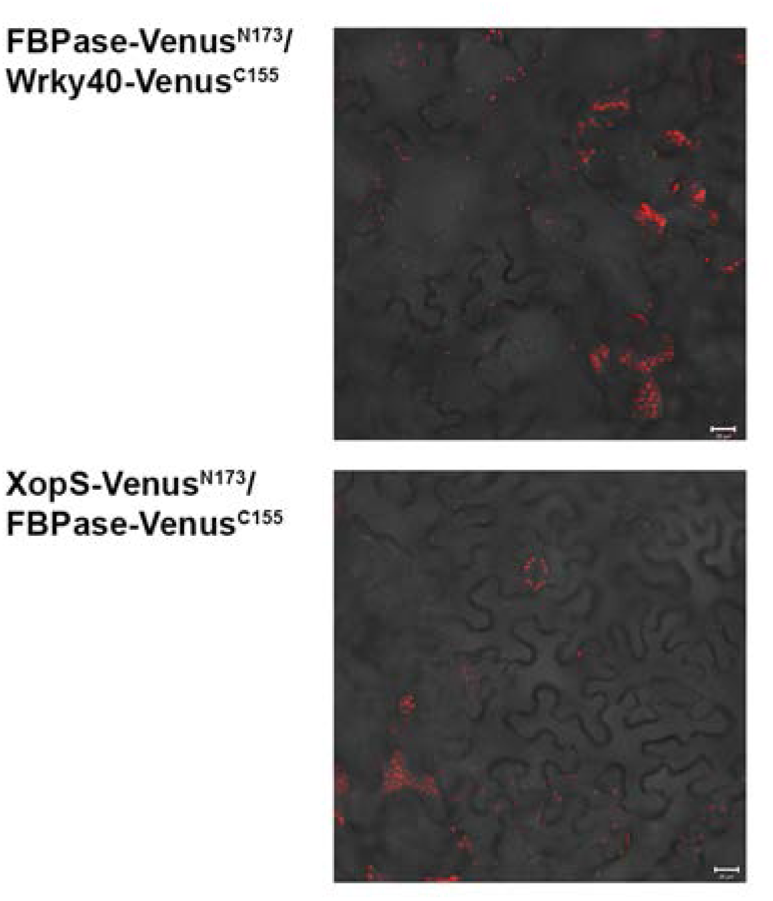
Control experiments for BiFC assays. YFP confocal microscopy images show a merge YFP- and chlorophyll auto-fluorescence of *N. benthamiana* leaf epidermal cells transiently expressing constructs encoding the fusion proteins indicated. Scale bars represent 20 µm. Each image is the representative of at least three experiments.

**Supplementary Figure S9.**
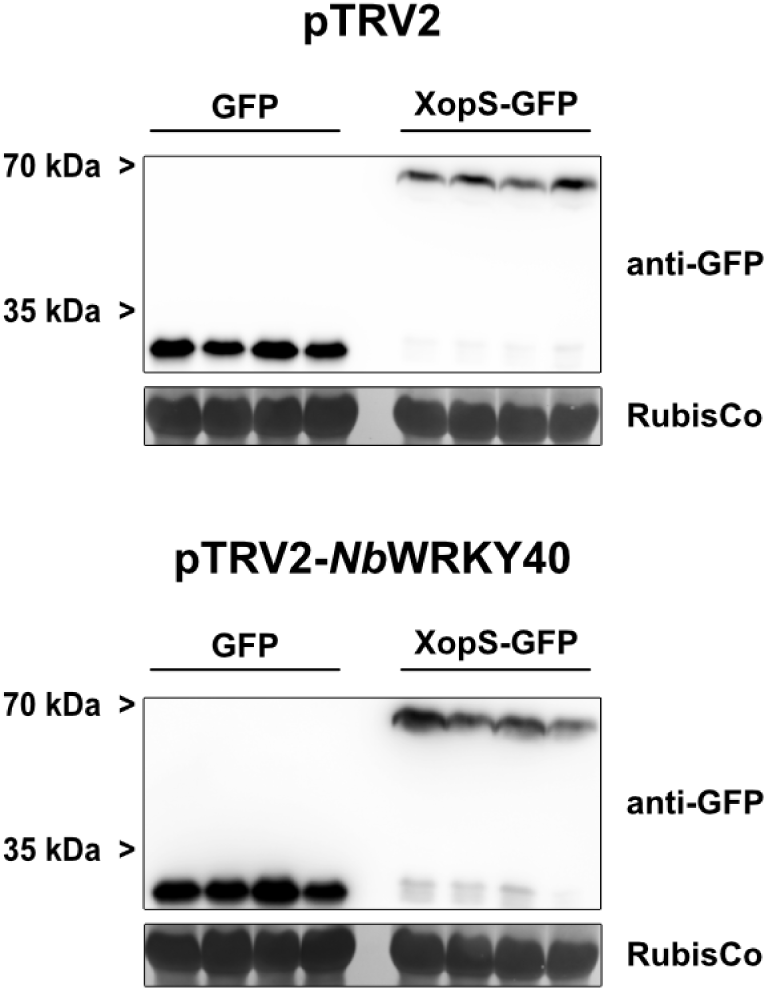
Verification of protein expression in pTRV2 (empty vector silencing; control) and pTRV2*-NbWRKY40* (*NbWRKY40* silencing) plants transiently expressing either GFP or XopS-GFP. Total protein extracts from *Agrobacterium*-infiltrated leaves were prepared 48 hpi and protein expression of indicated proteins was detected by western blotting using an anti-GFP antibody. Amido black staining of RubisCo served as loading control. The four biological replicates used for the analysis of stomatal aperture are shown.

**Supplementary Figure S10.**
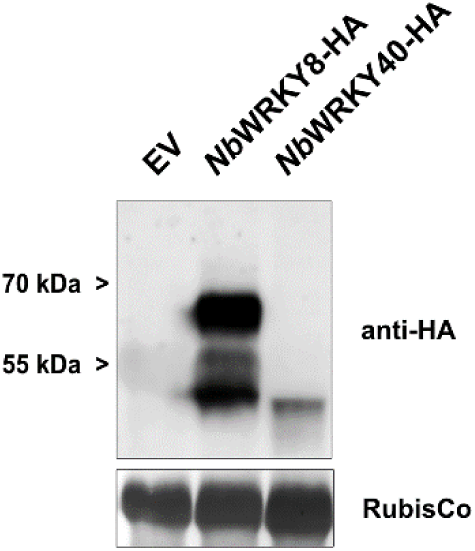
Verification of effector protein expression in *W-box::GUS* reporter gene analyses. Total protein extracts from *Agrobacterium*-infiltrated leaves were prepared 48 hpi and protein expression was immunodetected using an anti-HA antibody. Amido black staining of RubisCo served as loading control. Leaf discs from the biological replicates used for *W-box::GUS* reporter gene analyses were pooled for the western blot analysis shown here.

**Supplementary Figure S11.**
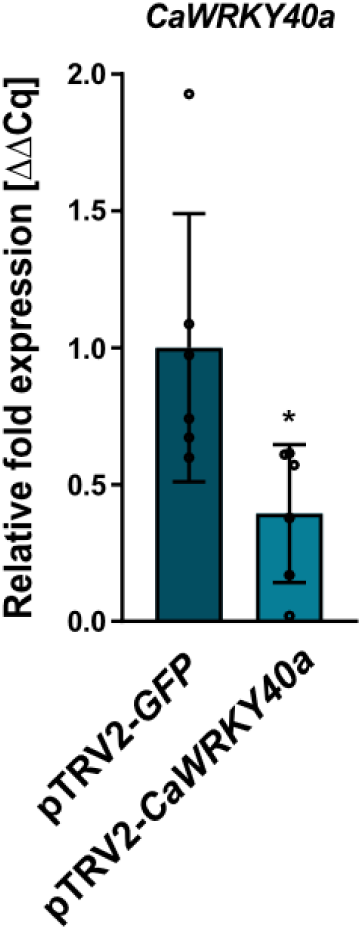
Verification of *CaWRKY40a* down-regulation in pTRV2-*CaWRKY40a* (*CaWRKY40a* silencing) compared to pTRV2-*GFP* (*GFP* silencing; control) in virus-induced-gene-silencing (VIGS) pepper plants used for defense gene expression analysis. Three weeks after infiltrating pepper plants with the silencing constructs, total RNA was isolated from excised leaves treated with 5mM SA for 4 h. The mRNA level of *CaWRKY40a* in pTRV2-*CaWRK40a* was measured by qRT-PCR and compared to *CaWRKY40a* expression in pTRV2-*GFP* control plants. *Ubiquitin* was used as a reference gene. Bars represent the mean of n = 6 biological replicates ± SD. Asterisks (*, P < 0.05) mark significant differences according to Student’s *t*-test.

**Supplementary Figure S12.**
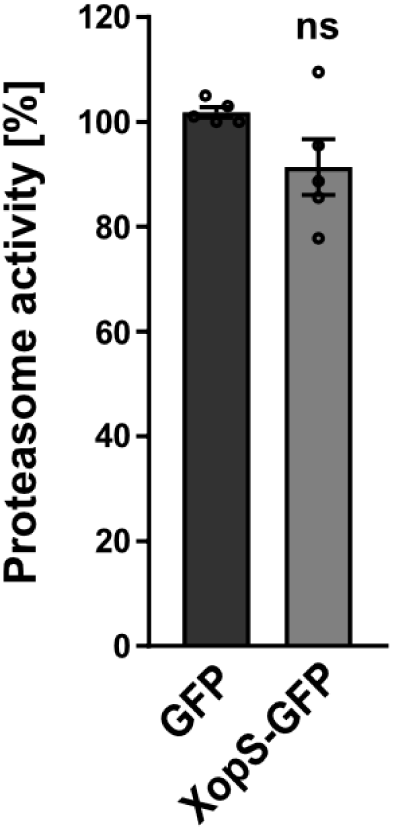
Expression of XopS does not affect proteasome activity. Proteasome activity is shown in *N. benthamiana* leaves following transient expression of XopS or GFP control. Relative proteasome activity in total protein extracts was determined by monitoring the breakdown of the fluorogenic peptide Suc-LLVY-AMC at 30°C in a fluorescence spectrophotometer. The GFP control was set to 100%. Bars represent the mean of n = 5 biological replicates ± SE. ns, not significant according to Student’s *t*-test.

**Supplementary Figure S13.**
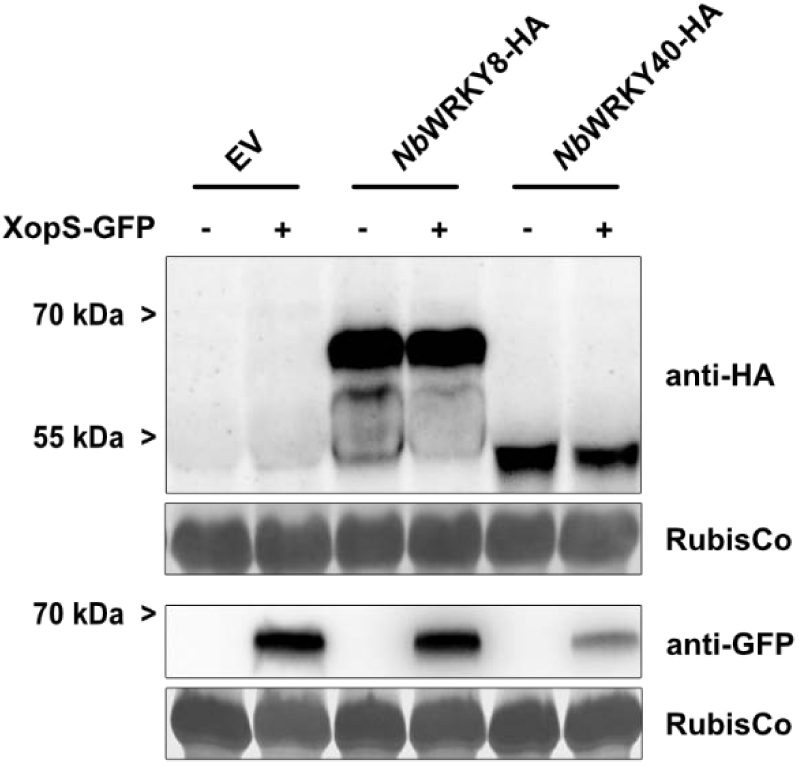
Verification of effector protein expression in *pCaPR4::GUS* reporter gene analyses. Total protein extracts from *Agrobacterium*-infiltrated leaves were prepared 48 hpi and protein expression was immune detected using an anti-HA or anti-GFP antibody. Amido black staining of RubisCo served as loading control. Leaf discs from the biological replicates used for *pCaPR4::GUS* reporter gene analyses were pooled for the western blot analysis shown here.

**Supplementary Figure S14.**
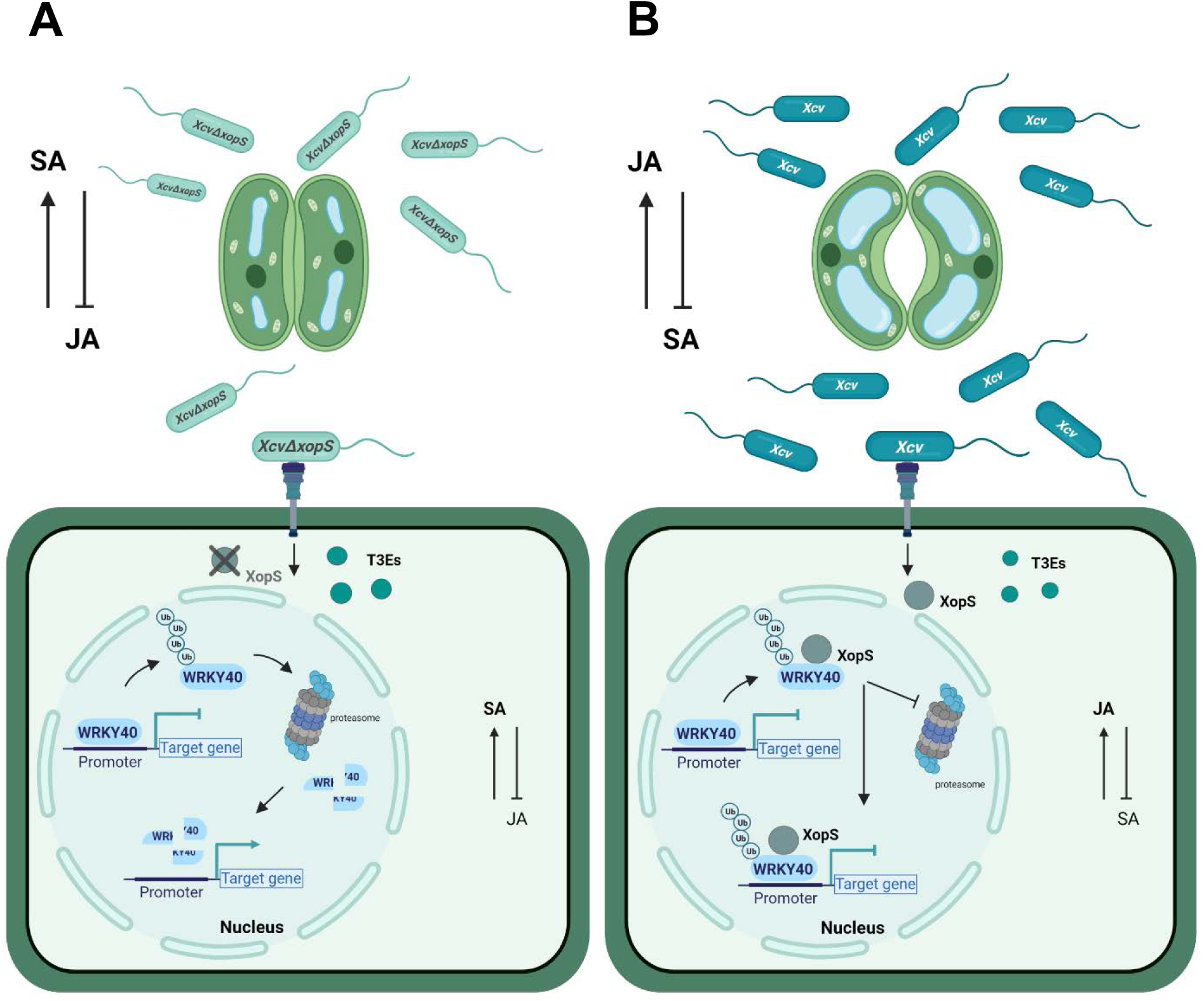
Working model of how XopS promotes *Xcv* virulence. **(A)** Upon infection of pepper plants with an *Xcv* strain lacking XopS, WRKY40 ubiquitination results in its degradation by the proteasome and leads to the de-repression of its target genes. This enables the induction of SA-dependent gene expression, such as *PR1*, and at the same time represses JA-dependent responses through increased expression of negative regulators, including *JAZ8*. As a result, stomata close in response to MAMP perception and the plant mounts apoplastic defense responses. **(B)** In contrast, wild type *Xcv* delivers XopS into the host cell, which prevents degradation of WRKY40 through physical interaction. Stabilized WRKY40 attenuates the induction of SA-dependent immunity and also mediates decreased expression of *JAZ8*. The latter results in an activation of JA-responses further antagonizing SA-mediated defense, including interference with stomatal closure to facilitate bacterial tissue entry.

**Supplementary Table 1.**
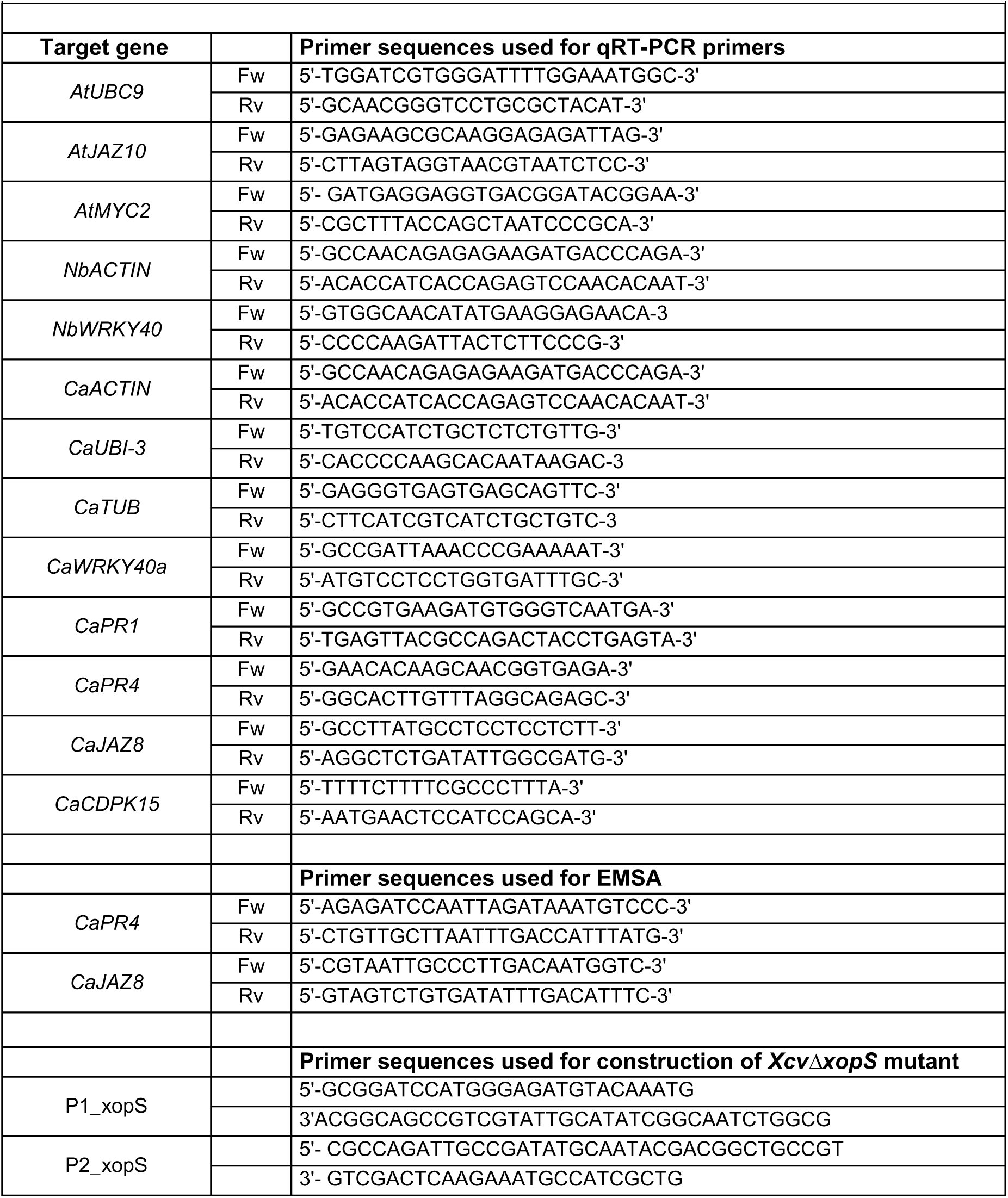
Nucleotide sequences of gene-specific primers.

## References

Arsova, B., Hoja, U., Wimmelbacher, M., Greiner, E., Üstün, S., Melzer, M., Petersen, K., Lein, W., and Börnke, F. (2010). Plastidial thioredoxin z interacts with two fructokinase-like proteins in a thiol-dependent manner: evidence for an essential role in chloroplast development in Arabidopsis and *Nicotiana benthamiana*. Plant Cell 22, 1498–1515.

Bakshi, M., and Oelmüller, R. (2014). WRKY transcription factors. Plant Signal Behav 9, e27700.

Birkenbihl, R.P., Kracher, B., Roccaro, M., and Somssich, I.E. (2017). Induced genome-wide binding of three Arabidopsis WRKY transcription factors during early MAMP-triggered immunity. Plant Cell 29, 20–38.

Bleckmann, A., Weidtkamp-Peters, S., Seidel, C.A.M., and Simon, R. (2010). Stem cell signaling in Arabidopsis requires CRN to localize CLV2 to the plasma membrane. Plant Physiol 152, 166–176.

Boch, J., and Bonas, U. (2010). Xanthomonas AvrBs3 family-type III effectors: discovery and function. Ann Rev Phytopathol 48, 419–436.

Boller, T., and Felix, G. (2009). A renaissance of elicitors: perception of microbe-associated molecular patterns and danger signals by pattern-recognition receptors. Annu Rev Plant Biol 60, 379–406.

Börnke, F. (2005). The variable C-terminus of 14-3-3 proteins mediates isoform-specific interaction with sucrose-phosphate synthase in the yeast two-hybrid system. J Plant Physiol 162, 161–168.

Büttner, D. (2016). Behind the lines-actions of bacterial type III effector proteins in plant cells. FEMS Microbiol Rev 40, 894–937.

Cesari, S. (2018). Multiple strategies for pathogen perception by plant immune receptors. New Phytol 219, 17–24.

Chen, H., Chen, J., Li, M., Chang, M., Xu, K., Shang, Z., Zhao, Y., Palmer, I., Zhang, Y., McGill, J., Alfano, J.R., Nishimura, M.T., Liu, F., and Fu, Z.Q. (2017). A bacterial type III effector targets the master regulator of salicylic acid signaling, NPR1, to subvert plant immunity. Cell Host Microbe 22, 777–788.e777.

Chi, Y., Yang, Y., Zhou, Y., Zhou, J., Fan, B., Yu, J.-Q., and Chen, Z. (2013). Protein–protein interactions in the regulation of WRKY transcription factors. Mol Plant 6, 287–300.

Clough, S.J., and Bent, A.F. (1998). Floral dip: a simplified method for Agrobacterium-mediated transformation of *Arabidopsis thaliana*. Plant J 16, 735–743.

Creelman, R.A., and Mullet, J.E. (1995). Jasmonic acid distribution and action in plants: regulation during development and response to biotic and abiotic stress. Proc Natl Acad Sci U S A 92, 4114–4119.

Dang, F.-F., Wang, Y.N., Yu, L., Eulgem, T., Lai, Y., Liu, Z.Q., Wang, X., Qiu, A.L., Zhang, T.X., Lin, J., Chem, Y.S., Guan, D.Y., Cai, H.Y., Mou, S.L., and He, S.L. (2013). CaWRKY40, a WRKY protein of pepper, plays an important role in the regulation of tolerance to heat stress and resistance to Ralstonia solanacearum infection. Plant Cell Environ 36, 757–774.

Dodds, P.N., and Rathjen, J.P. (2010). Plant immunity: towards an integrated view of plant-pathogen interactions. Nat Rev Genet 11, 539–548.

Dou, D., and Zhou, J.-M. (2012). Phytopathogen effectors subverting host immunity: different foes, similar battleground. Cell Host Microbe 12, 484–495.

Du, M., Zhai, Q., Deng, L., Li, S., Li, H., Yan, L., Huang, Z., Wang, B., Jiang, H., Huang, T., Li, C.- B., Wei, J., Kang, L., Li, J., and Li, C. (2014). Closely related NAC transcription factors of tomato differentially regulate stomatal closure and reopening during pathogen attack. Plant Cell 26, 3167–3184.

Eulgem, T., Rushton, P.J., Robatzek, S., and Somssich, I.E. (2000). The WRKY superfamily of plant transcription factors. Trends Plant Sci 5, 199–206.

Finley, D. (2009). Recognition and processing of ubiquitin-protein conjugates by the proteasome. Annu Rev Biochem 78, 477–513.

Garner, C.M., Kim, S.H., Spears, B.J., and Gassmann, W. (2016). Express yourself: Transcriptional regulation of plant innate immunity. Sem Cell Devel Biol 56, 150–162.

Gimenez-Ibanez, S., Hann, D.R., Ntoukakis, V., Petutschnig, E., Lipka, V., and Rathjen, J.P. (2009). AvrPtoB targets the LysM receptor kinase CERK1 to promote bacterial virulence on plants. Curr Biol 19, 423–429.

Gimenez-Ibanez, S., Boter, M., Fernández-Barbero, G., Chini, A., Rathjen, J.P., and Solano, R. (2014). The bacterial effector HopX1 targets JAZ transcriptional repressors to activate jasmonate signaling and promote infection in Arabidopsis. Plos Biol 12, e1001792.

Gimenez-Ibanez, S., Boter, M., Ortigosa, A., García-Casado, G., Chini, A., Lewsey, M.G., Ecker, J.R., Ntoukakis, V., and Solano, R. (2017). JAZ2 controls stomata dynamics during bacterial invasion. New Phytol 213, 1378–1392.

Göhre, V., Spallek, T., Haweker, H., Mersmann, S., Mentzel, T., Boller, T., de Torres, M., Mansfield, J.W., and Robatzek, S. (2008). Plant pattern-recognition receptor FLS2 is directed for degradation by the bacterial ubiquitin ligase AvrPtoB. Curr Biol 18, 1824–1832.

Higashi, K., Ishiga, Y., Inagaki, Y., Toyoda, K., Shiraishi, T., and Ichinose, Y. (2008). Modulation of defense signal transduction by flagellin-induced WRKY41 transcription factor in *Arabidopsis thaliana*. Mol Genet Genom 279, 303–312.

Huh, S.U., Lee, G.-J., Jung, J.H., Kim, Y., Kim, Y.J., and Paek, K.-H. (2015). *Capsicum annuum* transcription factor WRKYa positively regulates defense response upon TMV infection and is a substrate of CaMK1 and CaMK2. Sci Rep 5, 7981.

Hurley, B., Lee, D., Mott, A., Wilton, M., Liu, J., Liu, Y.C., Angers, S., Coaker, G., Guttman, D.S., and Desveaux, D. (2014). The *Pseudomonas syringae* type III effector HopF2 suppresses Arabidopsis stomatal immunity. PLoS One 9, e114921.

Ishihama, N., and Yoshioka, H. (2012). Post-translational regulation of WRKY transcription factors in plant immunity. Curr Opin Plant Biol 15, 431–437.

Ishihama, N., Yamada, R., Yoshioka, M., Katou, S., and Yoshioka, H. (2011). Phosphorylation of the *Nicotiana benthamiana* WRKY8 transcription factor by MAPK functions in the defense response. Plant Cell 23, 1153–1170.

Jones, J.B., Stall, R.E., and Bouzar, H. (1998). Diversity among xanthomonads pathogenic on pepper and tomato. Annu Rev Phytopathol 36, 41–58.

Karimi, M., Inzé, D., and Depicker, A. (2002). GATEWAY^(TM)^ vectors for *Agrobacterium*-mediated plant transformation. Trends Plant Sci 7, 193–195.

Katsir, L., Schilmiller, A.L., Staswick, P.E., He, S.Y., and Howe, G.A. (2008). COI1 is a critical component of a receptor for jasmonate and the bacterial virulence factor coronatine. Proc Natl Acad Sci 105, 7100–7105.

Kay, S., Hahn, S., Marois, E., Hause, G., and Bonas, U. (2007). A bacterial effector acts as a plant transcription factor and induces a cell size regulator. Science 318, 648–651.

Khan, M., Seto, D., Subramaniam, R., and Desveaux, D. (2018). Oh, the places they’ll go! A survey of phytopathogen effectors and their host targets. Plant J 93, 651–663.

Kim, J.G., Stork, W., and Mudgett, M.B. (2013). *Xanthomonas* type III effector XopD desumoylates tomato transcription factor SlERF4 to suppress ethylene responses and promote pathogen growth. Cell Host Microbe 13, 143–154.

Kim, J.G., Taylor, K.W., Hotson, A., Keegan, M., Schmelz, E.A., and Mudgett, M.B. (2008). XopD SUMO protease affects host transcription, promotes pathogen growth, and delays symptom development in xanthomonas-infected tomato leaves. Plant Cell 20, 1915–1929.

Krumm, T., Bandemer, K., and Boland, W. (1995). Induction of volatile biosynthesis in the lima bean (*Phaseolus lunatus*) by leucine- and isoleucine conjugates of 1-oxo- and 1-hydroxyindan-4- carboxylic acid: evidence for amino acid conjugates of jasmonic acid as intermediates in the octadecanoid signalling pathway. FEBS Lett 377, 523–529.

Kulathu, Y., and Komander, D. (2012). Atypical ubiquitylation - the unexplored world of polyubiquitin beyond Lys48 and Lys63 linkages. Nat Rev Mol Cell Biol 13, 508–523.

Langin, G., Gouguet, P., and Ustun, S. (2020). Microbial Effector Proteins - A Journey through the Proteolytic Landscape. Trends Microbiol 28, 523–535.

Leong, J.X., Raffeiner, M., Spinti, D., Langin, G., Franz-Wachtel, M., Guzman, A.R., Kim, J.-G., Pandey, P., Minina, A.E., Macek, B., Hafrén, A., Bozkurt, T.O., Mudgett, M.B., Börnke, F., Hofius, D., and Üstün, S. (2021). Self-ubiquitination of a pathogen type-III effector traps and blocks the autophagy machinery to promote disease. bioRxiv, 2021.2003.2017.435853.

Le Roux, C., Huet, G., Jauneau, A., Camborde, L., Trémousaygue, D., Kraut, A., Zhou, B., Levaillant, M., Adachi, H., Yoshioka, H., Raffaele, S., Berthomé, R., Couté, Y., Parker, J.E., and Deslandes, L. (2015). A receptor pair with an integrated decoy converts pathogen disabling of transcription factors to immunity. Cell 161, 1074–1088.

Lewis, L.A., Polanski, K., de Torres-Zabala, M., Jayaraman, S., Bowden, L., Moore, J., Penfold, C.A., Jenkins, D.J., Hill, C., Baxter, L., Kulasekaran, S., Truman, W., Littlejohn, G., Prusinska, J., Mead, A., Steinbrenner, J., Hickman, R., Rand, D., Wild, D.L., Ott, S., Buchanan-Wollaston, V., Smirnoff, N., Beynon, J., Denby, K., and Grant, M. (2015). Transcriptional dynamics driving MAMP-triggered immunity and pathogen effector-mediated immunosuppression in Arabidopsis leaves following infection with *Pseudomonas syringae* pv *tomato* DC3000. Plant Cell 27, 3038–3064.

Li, B., Meng, X., Shan, L., and He, P. (2016). Transcriptional regulation of pattern-triggered immunity in plants. Cell Host Microbe 19, 641–650.

Lichtenthaler, H.K. (1987). Chlorophylls and carotenoids, pigments of photosynthetic biomembranes. Meth Enzymol 148, 350–382.

Lin, Y., Hu, Q., Zhou, J., Yin, W., Yao, D., Shao, Y., Zhao, Y., Guo, B., Xia, Y., Chen, Q., Wang, Y., Ye, W., Xie, Q., Tyler, B.M., Xing, W., and Wang, Y. (2021). *Phytophthora sojae* effector Avr1d functions as an E2 competitor and inhibits ubiquitination activity of GmPUB13 to facilitate infection. Proc Natl Acad Sci U S A 118.

Liu, B., Jiang, Y., Tang, H., Tong, S., Lou, S., Shao, C., Zhang, J., Song, Y., Chen, N., Bi, H., Zhang, H., Li, J., Liu, J., and Liu, H. (2021). The ubiquitin E3 ligase SR1 modulates the dubmergence response by degrading phosphorylated WRKY33 in Arabidopsis. Plant Cell.

Liu, Y., Schiff, M., and Dinesh-Kumar, S.P. (2002). Virus-induced gene silencing in tomato. Plant J 31, 777–786.

Lozano-Duran, R., Bourdais, G., He, S.Y., and Robatzek, S. (2014). The bacterial effector HopM1 suppresses PAMP-triggered oxidative burst and stomatal immunity. New Phytol 202, 259–269.

Macho, A.P., and Zipfel, C. (2015). Targeting of plant pattern recognition receptor-triggered immunity by bacterial type-III secretion system effectors. Curr Opin Microbiol 23C, 14–22.

Mao, G., Meng, X., Liu, Y., Zheng, Z., Chen, Z., and Zhang, S. (2011). Phosphorylation of a WRKY transcription factor by two pathogen-responsive MAPKs drives phytoalexin biosynthesis in Arabidopsis. Plant Cell 23, 1639–1653.

Matsushita, A., Inoue, H., Goto, S., Nakayama, A., Sugano, S., Hayashi, N., and Takatsuji, H. (2013). Nuclear ubiquitin proteasome degradation affects WRKY45 function in the rice defense program. Plant J 73, 302–313.

McGuire, R.G., Jones, J.B., and Scott, J.W. (1991). Epiphytic populations of *Xanthomonas campestris* pv. *vesicatoria* on tomato cultigens resistant and susceptible to bacterial spot. Plant Disease 75, 606–609.

Melotto, M., Underwood, W., and He, S.Y. (2008). Role of stomata in plant innate immunity and foliar bacterial diseases. Annu Rev Phytopathol 46, 101–122.

Melotto, M., Underwood, W., Koczan, J., Nomura, K., and He, S.Y. (2006). Plant stomata function in innate immunity against bacterial invasion. Cell 126, 969–980.

Miao, Y., and Zentgraf, U. (2010). A HECT E3 ubiquitin ligase negatively regulates Arabidopsis leaf senescence through degradation of the transcription factor WRKY53. Plant J 63, 179–188.

Mittal, S., and Davis, K.R. (1995). Role of the phytotoxin coronatine in the infection of Arabidopsis thaliana by *Pseudomonas syringae* pv. *tomato*. Mol Plant Microbe Interact 8, 165–171.

Moore, J.W., Loake, G.J., and Spoel, S.H. (2011). Transcription dynamics in plant immunity. Plant Cell 23, 2809–2820.

Mukhopadhyay, D., and Riezman, H. (2007). Proteasome-independent functions of ubiquitin in endocytosis and signaling. Science 315, 201–205.

Nakamura, S., Mano, S., Tanaka, Y., Ohnishi, M., Nakamori, C., Araki, M., Niwa, T., Nishimura, M., Kaminaka, H., Nakagawa, T., Sato, Y., and Ishiguro, S. (2010). Gateway binary vectors with the bialaphos resistance gene, bar, as a selection marker for plant transformation. Biosci Biotechnol Biochem 74, 1315–1319.

Nietzsche, M., Guerra, T., Alseekh, S., Wiermer, M., Sonnewald, S., Fernie, A.R., and Börnke, F. (2018). STOREKEEPER RELATED1/G-element binding protein (STKR1) interacts with protein kinase SnRK1. Plant Physiol 176, 1773–1792.

Oh, S.K., Baek, K.H., Park, J.M., Yi, S.Y., Yu, S.H., Kamoun, S., and Choi, D. (2008). *Capsicum annuum* WRKY protein CaWRKY1 is a negative regulator of pathogen defense. New Phytol 177, 977–989.

Okada, M., Ito, S., Matsubara, A., Iwakura, I., Egoshi, S., and Ueda, M. (2009). Total syntheses of coronatines by exo-selective Diels–Alder reaction and their biological activities on stomatal opening. Organ Biomol Chem 7, 3065–3073.

Pandey, S.P., and Somssich, I.E. (2009). The role of WRKY transcription factors in plant immunity. Plant Physiol 150, 1648–1655.

Pandey, S.P., Roccaro, M., Schon, M., Logemann, E., and Somssich, I.E. (2010). Transcriptional reprogramming regulated by WRKY18 and WRKY40 facilitates powdery mildew infection of Arabidopsis. Plant J 64, 912–923.

Popov, G., Fraiture, M., Brunner, F., and Sessa, G. (2016). Multiple *Xanthomonas euvesicatoria* type III effectors inhibit flg22-triggered immunity. Mol Plant Microbe Interact 29, 651–660.

Potnis, N., Krasileva, K., Chow, V., Almeida, N.F., Patil, P.B., Ryan, R.P., Sharlach, M., Behlau, F., Dow, J.M., Momol, M.T., White, F.F., Preston, J.F., Vinatzer, B.A., Koebnik, R., Setubal, J.C., Norman, D.J., Staskawicz, B.J., and Jones, J.B. (2011). Comparative genomics reveals diversity among xanthomonads infecting tomato and pepper. BMC Genomics 12, 146.

Ramos, L.J., and Volin, R.B. (1987). Role of stomatal opening and frequency on infection of *Lycopersicon* spp. by *Xanthomonas campestris* pv. *vesicatoria*. Phytopathol 77, 1311–1317.

Rosli, H.G., Zheng, Y., Pombo, M.A., Zhong, S., Bombarely, A., Fei, Z., Collmer, A., and Martin, G.B. (2013). Transcriptomics-based screen for genes induced by flagellin and repressed by pathogen effectors identifies a cell wall-associated kinase involved in plant immunity. Genome Biol 14, R139.

Rushton, P.J., Somssich, I.E., Ringler, P., and Shen, Q.J. (2010). WRKY transcription factors. Trends Plant Sci 15, 247–258.

Sarris, P.F., Duxbury, Z., Huh, S.U., Ma, Y., Segonzac, C., Sklenar, J., Derbyshire, P., Cevik, V., Rallapalli, G., Saucet, S.B., Wirthmueller, L., Menke, F.L.H., Sohn, K.H., and Jones, J.D.G. (2015). A plant immune receptor detects pathogen effectors that target WRKY transcription factors. Cell 161, 1089–1100.

Sawinski, K., Mersmann, S., Robatzek, S., and Böhmer, M. (2013). Guarding the green: pathways to stomatal immunity. Mol Plant Microbe Interact 26, 626–632.

Schulze, S., Kay, S., Büttner, D., Egler, M., Eschen-Lippold, L., Hause, G., Krüger, A., Lee, J., Müller, O., Scheel, D., Szczesny, R., Thieme, F., and Bonas, U. (2012). Analysis of new type III effectors from Xanthomonas uncovers XopB and XopS as suppressors of plant immunity. New Phytol 195, 894–911.

Schwartz, A.R., Potnis, N., Timilsina, S., Wilson, M., Patané, J., Martins, J., Minsavage, G.V., Dahlbeck, D., Akhunova, A., Almeida, N., Vallad, G.E., Barak, J.D., White, F.F., Miller, S.A., Ritchie, D., Goss, E., Bart, R.S., Setubal, J.C., Jones, J.B., and Staskawicz, B.J. (2015). Phylogenomics of *Xanthomonas* field strains infecting pepper and tomato reveals diversity in effector repertoires and identifies determinants of host specificity. Front Microbiol 6.

Shen, L., Yang, S., Yang, T., Liang, J., Cheng, W., Wen, J., Liu, Y., Li, J., Shi, L., Tang, Q., Shi, W., Hu, J., Liu, C., Zhang, Y., Mou, S., Liu, Z., Cai, H., He, L., Guan, D., Wu, Y., and He, S. (2016). CaCDPK15 positively regulates pepper responses to *Ralstonia solanacearum* inoculation and forms a positive-feedback loop with CaWRKY40 to amplify defense signaling. Sci Rep 6, 22439.

Staswick, P.E., and Tiryaki, I. (2004). The oxylipin signal jasmonic acid is activated by an enzyme that conjugates it to isoleucine in Arabidopsis. Plant Cell 16, 2117–2127.

Teper, D., Burstein, D., Salomon, D., Gershovitz, M., Pupko, T., and Sessa, G. (2016). Identification of novel *Xanthomonas euvesicatoria* type III effector proteins by a machine-learning approach. Mol Plant Pathol 17, 398–411.

Tsuda, K., and Katagiri, F. (2010). Comparing signaling mechanisms engaged in pattern-triggered and effector-triggered immunity. Curr Opin Plant Biol 13, 459–465.

Tsuda, K., and Somssich, I.E. (2015). Transcriptional networks in plant immunity. New Phytol 206, 932–947.

Üstün, S., and Börnke, F. (2017). Ubiquitin proteasome activity measurement in total plant extracts. Bio-protocol 7, e2532.

Üstün, S., Bartetzko, V., and Börnke, F. (2013). The *Xanthomonas campestris* type III effector XopJ targets the host cell proteasome to suppress salicylic-acid mediated plant defence. PLoS Pathog 9, e1003427.

Üstün, S., König, P., Guttman, D.S., and Börnke, F. (2014). HopZ4 from *Pseudomonas syringae*, a member of the HopZ type III effector family from the YopJ superfamily, inhibits the proteasome in plants. Mol Plant Microb Interact 27, 611–623.

Üstün, S., Sheikh, A., Gimenez-Ibanez, S., Jones, A., Ntoukakis, V., and Börnke, F. (2016). The proteasome acts as a hub for plant immunity and is targeted by Pseudomonas type III effectors. Plant Physiol 172, 1941–1958.

Vierstra, R.D. (2009). The ubiquitin-26S proteasome system at the nexus of plant biology. Nat Rev Mol Cell Biol 10, 385–397.

Xu, X., Chen, C., Fan, B., and Chen, Z. (2006). Physical and functional interactions between pathogen-induced Arabidopsis WRKY18, WRKY40, and WRKY60 transcription factors. Plant Cell 18, 1310–1326.

Ye, Q., Wang, H., Su, T., Wu, W.-H., and Chen, Y.-F. (2018). The ubiquitin E3 ligase PRU1 tegulates WRKY6 degradation to modulate phosphate homeostasis in response to low-Pi stress in Arabidopsis. Plant Cell 30, 1062–1076.

Ye, R., Yao, Q.-H., Xu, Z.-H., and Xue, H.-W. (2004). Development of an efficient method for the isolation of factors involved in gene transcription during rice embryo development. Plant J 38, 348–357.

Zhang, Y.X., Callaway, E.M., Jones, J.B., and Wilson, M. (2009). Visualisation of *hrp* gene expression in *Xanthomonas euvesicatoria* in the tomato phyllosphere. Eur J Plant Pathol 124, 379–390.

Zheng, X.Y., Spivey, N.W., Zeng, W., Liu, P.P., Fu, Z.Q., Klessig, D.F., He, S.Y., and Dong, X. (2012). Coronatine promotes *Pseudomonas syringae* virulence in plants by activating a signaling cascade that inhibits salicylic acid accumulation. Cell Host Microbe 11, 587–596.

Zipfel, C. (2014). Plant pattern-recognition receptors. Trends Immunol 35, 345–351.

